# Metabolic complexity in the human brain supports metastable network dynamics and declines in normative ageing

**DOI:** 10.64898/2026.11.02.763776

**Authors:** Hamish A. Deery, Emma X. Liang, Chris Moran, Gary F. Egan, Sharna D. Jamadar

## Abstract

The brain’s adaptive capacity is tied to the complexity of its neural activity. While entropy of brain signals has been used to characterise this complexity in haemodynamic and electrophysiological systems, it remains unknown whether entropy contributes to the temporal dynamics of the brain’s metabolic signals (glucodynamics) and how metabolic entropy may relate to network organisation, brain ageing and cognitive decline. Here, we combined dynamic functional PET (fPET) with ^18^F-fluoroedeoxyglucose and simultaneously acquired functional magnetic resonance imaging (fMRI) in 84 adults to investigate multiscale entropy (MSE) in the brain. We report that metabolic complexity measured with PET MSE represents non-linear, high-frequency complexity, whereas hemodynamic complexity reflects non-linearly constrained neurovascular phase dynamics. Greater metabolic complexity was associated with greater network metastability, particularly in unimodal and higher-order associative networks. Greater regional metabolic complexity also correlated with greater cortical thickness, cerebral metabolic rates of glucose (CMR_GLC_) and cerebral blood flow, and was lower in older than younger adults. In contrast, global and network haemodynamic MSE was not significant associated with network dynamics. Higher global metabolic complexity was associated with better cognitive performance, and voxel-level derived MSE clusters were \associated with cognitive performance in both modalities, with varied cluster sizes and spatial locations. However, these associations were largely accounted for by age variance. We conclude that temporal complexity in cerebral glucose is associated with the dynamics of the underlying metabolic signals, enables metastable network states and declines in ageing. Entropy of metabolic brain signals offers promise as a neurobiologically grounded metric of brain ageing.

## Introduction

The brain is a highly dynamic system, operating far from equilibrium and constantly reconfiguring its network architecture to meet fluctuating internal and external demands [1–4]. A central tenet of modern neuroscience is that this capacity for adaptation is intrinsically linked to the complexity of neural firing patterns [5]. Complexity, often operationalised as entropy, reflects the degree of unpredictability and information content within a system [5]. In natural and artificial systems, a balance between order and randomness is optimal; too much order (low entropy) results in rigidity and an inability to respond to novel stimuli, whereas too much randomness (high entropy) results in uncoordinated activity [6]. In the brain, this optimal complexity is thought to enable a flexible repertoire of network states necessary for adaptive cognition [7, 8]. Here, we explore whether the brain’s metabolic system exhibits temporal complexity measured by entropy of cerebral glucose metabolism.

Complexity is recognised as a defining feature of the large-scale network organisation of the brain. Functional network studies have established that *metastability* is a fundamental property of healthy brain function [9]. Metastability is the capacity of the brain to flexibly transition between network configurations without locking into a single state [9]. In the context of brain dynamics, ’state’ refers to the instantaneous level of phase synchrony among brain regions, with metastability measured as the temporal variability of this synchrony [2, 10]. Metastability is a core concept in the theory of Coordination Dynamics, which conceptualises the brain as a complex, self-organising system [2, 11, 12]. Within this framework, metastability reflects the dynamic balance between networks that synchronise (integrate) and function independently (segregate) to enable the local and global coordination necessary for cognitive processing [13, 14]. Underlying this metastable regime is the brain’s requirement for efficient information transfer, which is optimised when there is a balance between global integration and local specialisation [15, 16].

Large-scale brain dynamics are not an inconsequential by-product of rapid neural firing. Rather, they are intrinsically constrained by the brain’s metabolic system [17–20], which provides the energy needed for almost every aspect of brain function [21, 22]. Normative ageing and neurodegenerative diseases are accompanied by changes in brain metabolism [23–26]. Recent evidence suggests that the temporal dynamics of these metabolic signals, or ‘glucodynamics’ [17, 20], are not passive signals of local activity but carry biologically and functionally meaningful information. Regional glucodynamic signals covary over time, forming coherent large-scale networks that are separate but related to the brain’s fMRI-derived functional architecture and are differentially related to cognitive performance [27–31]. These temporally coherent metabolic signals exhibit a macroscale gradient organisation that aligns with cortical thickness, glucose metabolism (CMR_GLC_), cerebral blood flow (CBF) and gene expression [32]. These observations suggest that a highly organised, energy-dependent and temporally ordered architecture underlies brain function. However, it remains unknown whether the brain’s metabolic systems exhibit temporal complexity, and whether this complexity supports large scale networks to attain metastability.

To capture complexity of brain signals across timescales, multiscale entropy (MSE) has emerged as an important analytical tool [33, 34]. Unlike single-scale entropy measures, MSE quantifies signal complexity across a range of temporal scales, and thereby characterises a system’s information capacity across different timeframes [33]. MSE has been applied to electrophysiological (EEG) and fMRI data at the voxel and network-level, both at rest and during task conditions [35–38]. This work has revealed distinct MSE profiles across resting state functional brain networks [39–41] and reductions in entropy from rest to task states, including attention, executive control, learning and visual detection tasks [42, 43]. MSE is also altered in normative ageing [44, 45] and neurological and psychiatric conditions [46]. However, to our knowledge, the investigation of multiscale brain entropy has been confined to haemodynamic and electrophysiological signals, with the complexity of brain metabolism signals across timescales unknown.

Here, we address this gap. We leverage the unique temporal dynamics of functional FDG positron emission tomography (fPET) to investigate whether entropy contributes to the brain’s glucodynamic signals and how it may relate to metastable network dynamics, brain ageing and cognition. We compare fPET-derived metabolic complexity against simultaneously acquired fMRI data to determine whether these two modalities capture distinct or complementary information about brain complexity. Given that fPET and fMRI signals have different physiological bases [27, 29] and the connectivity of networks derived from those signals are differentially related to cognition [29, 31], we hypothesise that metabolic signals will exhibit MSE with distinct properties in comparison to fMRI (e.g., different mean and variability profiles). Second, we hypothesise that greater regional fPET MSE will be associated with greater network metastability. We also hypothesise that fPET MSE will spatially align to cortical thickness, baseline CMR_GLC_, and CBF. Finally, we hypothesise that fPET MSE will decline in normative ageing and that greater fPET MSE will be associated with better cognitive performance. By testing these hypotheses, our goal is to determine if fPET MSE can serve as a biologically grounded metric of brain complexity and provide a mechanistic link between metabolic signal dynamics, metastable network states, and age-related cognitive decline.

## Results

We investigated MSE in a sample of 84 healthy people who underwent a 90-minute, simultaneous fPET with ^18^F-flurodeoxyglucose (FDG) and fMRI scans while watching a naturalistic movie (see Supplement 1 for demographics). For each participant, MSE was calculated from 4D fPET (16s frames) and fMRI (TR = 1s volumes) images using a coarse-graining approach [33], with the following parameters: embedding dimension (m) = 2; tolerance threshold (r) = 0.15 × standard deviation of each voxel’s time series. For each scale factor (τ), the time series were coarse-grained by averaging consecutive data points in non-overlapping windows [33]. We initially explored a range of scaling factors: (τ) = 1, 2, 3 and 4 for fPET (16, 32, 48, 64 seconds) and 1, 2, 3, 4, 8, 16, 32 and 48 for fMRI (1-48 seconds). The larger number of time periods for fMRI were chosen to overlap with the temporal window of the fPET data. Based on qualitative and quantitative analysis of the MSE maps (reported below) and to allow for a degree of comparability between modalities, our primary analysis focussed on fPET periods of 16, 32, and 48 seconds (τ = 1, 2, 3) and fMRI periods of 4, 8 and 16 seconds (τ = 4, 8, 16). MSE was calculated using the Schaefer parcellation [47] at the100 region, 8 network and global (whole brain) levels.

To investigate the relationship between MSE and dynamic network topology, we computed a metastability index and an index of the balance between integration and segregation from the fPET and fMRI regional timeseries for each participant. We also calculated regional cortical thickness, CMR_GLC_, and CBF for each participant. For each modality and MSE period, we tested both linear and non-linear (quadratic) relationships across subjects between MSE, network metrics, cortical thickness, CMR_GLC,_ CBF and cognition. Age group differences (younger versus older) were also tested. Each analysis included head motion as a covariate and was corrected for multiple comparisons using false discovery rate corrections at p-FDR < 0.05.

We report analysis of the empirical MSE data against phase-randomised surrogates to aid interpretation of the biological meaning of the entropy measures. We then describe MSE within and across the fPET and fMRI modalities and report the associations between MSE, network properties and the other brain features. Finally, we report the age group differences in MSE and its association with cognition.

### MSE Signal Properties

The MSE of the fPET and fMRI timeseries is shown in Figure 1. To determine whether empirical MSE values capture non-linear dynamical complexity rather than linear autocorrelation or stochastic noise, we compared empirical MSE values against phase-randomised surrogate data (Figure 1A). Surrogate data serve as a statistical baseline model generated to match the linear properties of the empirical signal without its non-linear structure. Phase randomisation preserves the linear power spectral density and autocorrelation of the original time series while selectively destroying phase relationships and non-linear temporal dependencies.

**Figure 1.**
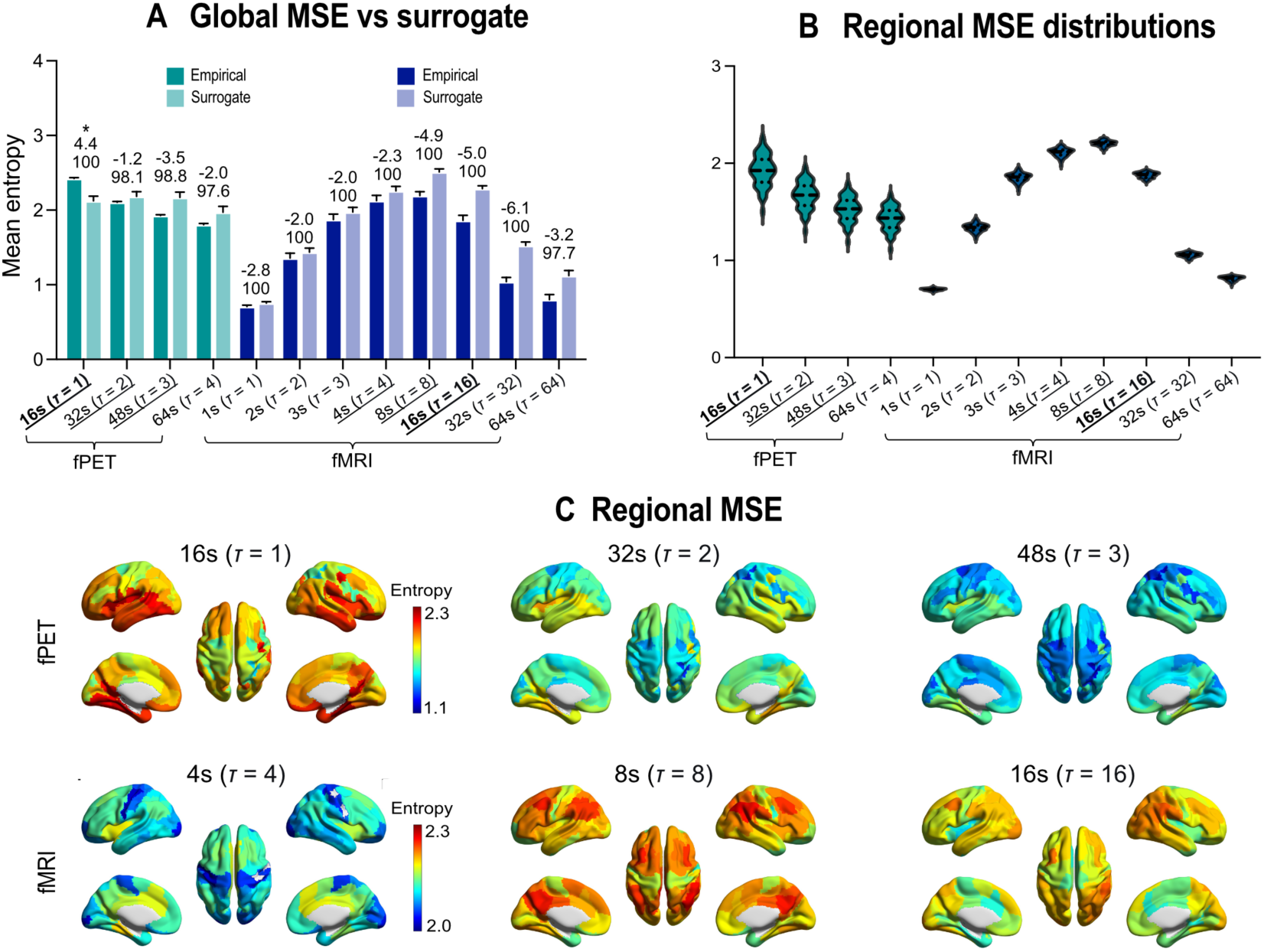
Multiscale entropy (MSE) descriptive statistics. (A) MSE was calculated at multiple temporal scales for fPET and fMRI and compared to surrogate null models. Bars show group-mean MSE for empirical and phase-randomised surrogate signals; error bars indicate SD across subjects. Values above each pair indicate Cohen’s d (paired, empirical - surrogate), with asterisk (*) denoting statistical significance (p-FDR < 0.05). Value below Cohen’s d are the percentage of subjects with |z| > 1.96 (i.e., empirical MSE falling outside the surrogate distribution). Only fPET MSE at 16s (*T* = 1) showed empirical values significantly above the surrogate null. Temporal periods of 16s, 32s and 48s (*T* = 1-3) in fPET and 4, 8, and 16s (*T* = 4, 8 and 16) in fMRI were retained for downstream analysis based on MSE values and to allow some degree of comparability across temporal scales. Those scales are underlined in panels (A), with bold font further indicating temporal correspondence across fPET and fMRI MSE (16s). (B) Regional MSE distributions for the 100 Schaefer regions, and (C) mean regional values projected to the brain surface.

For fPET at 16s (τ = 1), empirical MSE significantly exceeded surrogate values (mean empirical = 2.42, surrogate = 2.13), with 100% of subjects showing this effect (mean z = 55.1). These results confirm that fPET MSE at this fine temporal scale captures non-linear, high-frequency complexity characterised by high intrinsic signal irregularity (see Supplementary Figure 1 for illustration). Given that FDG-PET measures glucose use directly at the post-synaptic neuron [48–50], this elevation in empirical complexity relative to surrogates demonstrates that fPET signals encode rich, non-linear metabolic dynamics.

Conversely, for fMRI across all evaluated temporal scales (1-48s), empirical MSE values were significantly lower than those of their phase-randomised surrogates in 98-100% of participants (mean z = -120.3 at 1s to z = -11.7 at 48s; Figure 1A). In non-linear timeseries analysis, empirical entropy falling below phase-randomised surrogates is an established indicator of deterministic non-linear phase coupling and temporal structure [33, 51, 52]. Because phase-randomised surrogates preserve the linear power spectrum, and thus account for the smooth linear autocorrelation introduced by the haemodynamic response function (HRF), the lower entropy in empirical BOLD signal compared to surrogates confirms that haemodynamics operate as a relatively ordered, non-linearly constrained neurovascular system [53, 54].

Rather than reflecting statistical artifacts, these contrasting surrogate patterns demonstrate that fPET and fMRI MSE measure fundamentally distinct physiological features: fPET captures high-frequency non-linear metabolic fluctuations, whereas fMRI reflects non-linearly constrained, low-frequency haemodynamic fluctuations and deterministic temporal transitions. Consequently, both modalities provide complementary windows into brain signal complexity, with fPET MSE measuring high intrinsic signal irregularity and fMRI MSE capturing non-linear neurovascular regulation.

### Modality Differences in Regional MSE Characteristics

To test our hypothesis that metabolic signals exhibit complexity, albeit with distinct properties compared to fMRI (e.g., different mean and variability), we examined regional MSE across temporal scales. MSE values exhibited clear, modality-specific patterns (Figure 1B). For fPET, regional MSE values were highest at 16s (τ = 1) and monotonically decreased with increasing window length. In contrast, fMRI MSE followed a distinct inverted-U pattern, starting at low values at 1s (τ = 1), rising to peak at 8s (τ = 8), and then declining at longer window lengths. A within-subject general linear model comparing global fPET and fMRI MSE at the equivalent temporal scale of 16s (fPET τ = 1 vs fMRI τ = 16) revealed a significant difference between modalities (t(83) = 5.94, p < 0.001; Cohen’s d = 0.65), with fPET MSE significantly higher than fMRI MSE. Together, these results indicate that metabolic complexity is maximal and significantly higher at the native frame rate than haemodynamic complexity. In contrast, haemodynamic complexity peaks at intermediate timescales, consistent with previous fMRI reports [39, 41].

Regionally, both modalities exhibited a consistent spatial pattern, but with different network emphasis (Figure 1C; also see Supplement 4). For fPET, the highest MSE values were observed in unimodal sensory and somatomotor regions at 16s, notably the Somatomotor network (region B S2, MSE=2.18), the Visual network (peripheral region, MSE=2.16), and the temporal parietal junction (MSE=2.15). The lowest fPET MSE values were found in the intraparietal sulcus (IPS) of the Control network (LH: MSE=1.65; RH: MSE=1.49) and the frontal eye fields (FEF) of the Dorsal Attention network (MSE=1.64). For fMRI, the highest MSE values were found at 8s and localised to the insula of the Salience Ventral Attention network (MSE=2.19) and the parahippocampal cortex of the Default network (MSE=2.20). Conversely, the lowest fMRI MSE values were observed at 8s in the Somatomotor network (central region 1, MSE=2.13) and the temporal parietal junction (MSE=2.22). These results reveal a unimodal-to-association gradient, with fPET entropy peaking in sensorimotor cortex and fMRI entropy peaking in salience and default mode regions.

Based on these statistical properties, we retained MSE at temporal windows of 16, 32 and 48s for fPET (τ = 1, 2 and 3) and 4, 8 and 16s for fMRI (τ = 4, 8 and 16) in subsequent analyses. This selection optimises two key criteria: physiological validity and a degree of cross-modal temporal comparability. For fPET, 16-48s capture the window of maximal entropy. For fMRI, 4-16s capture the peak of the inverted-U trajectory. These ranges also provide a common timescale (16s) to compare signal complexity across modalities (effective timescales of 16-48s for fPET vs 4-16s for fMRI).

### MSE and Dynamic Network Features

Next, we used partial correlation analyses to test our hypothesis that greater fPET MSE will be associated with increased network metastability. A metastability index was computed as the standard deviation of global synchrony over time (see Methods for details). An integration-segregation balance index (ISBI) was calculated by dividing global efficiency by local efficiency (Figure 2A). ISBI values near 0.5 indicate an optimal network configuration, while deviations suggest a predisposition toward either integration (ISBI > 0.5) or segregation (ISBI < 0.5) [13].

**Figure 2.**
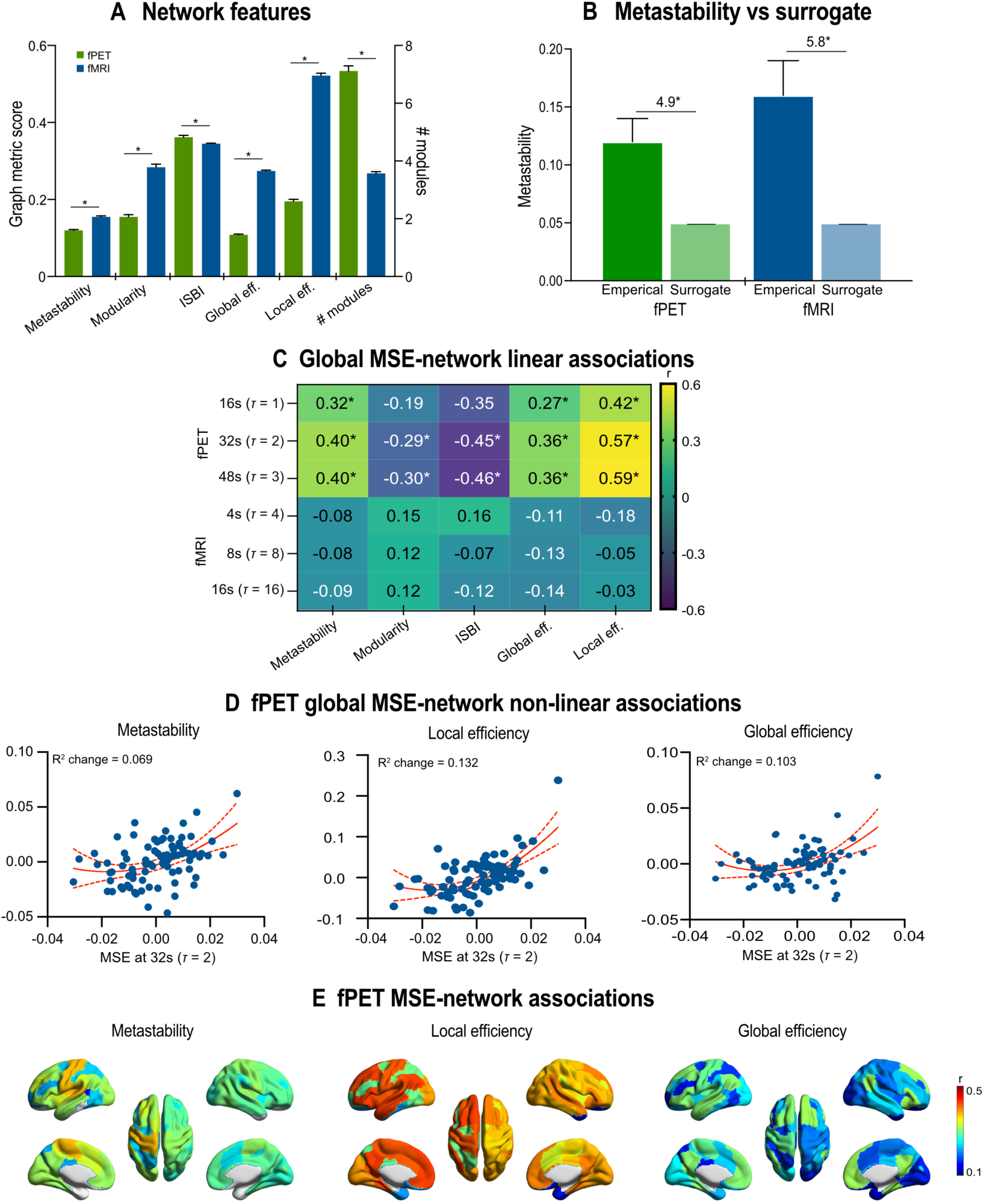
Multiscale entropy (MSE) relationships with network dynamics. (A) Mean (SE error bars) of network metrics including metastability, global efficiency, local efficiency, modularity, and the integration-segregation balance index (ISBI). fPET showed significantly higher modularity and number of modules, while fMRI showed higher absolute global and local efficiency. The ISBI indicated that fPET was organised for a closer-to-optimal integration-segregation balance (mean ISBI = 0.362 vs. 0.345, p = 0.002). (B) metastability index compared to surrogate null. Numbers above the bars are Cohen’s d. The empirical values was significantly higher than the surrogate (p < 0.001), indicating metastability captures genuine coordinated network dynamics in both fPET and fMRI. (C) Correlation analyses between MSE and network metrics. Heatmap shows partial correlation coefficients (r) with asterisk (*) denoting FDR-corrected significance (p-FDR < 0.05). Significant associations were found for fPET but not fMRI. (D) Significant non-linear (quadratic) relationships between fPET MSE and network metrics at 32s and 48s (*T* = 2 and 3), illustrating significant “J-shaped” relationships for metastability, global efficiency, and local efficiency (p-FDR < 0.05), in which the highest entropy values were associated with the highest network feature values. (E) Associations between network-level fPET MSE at 16s (*T* = 1) and network metrics. fMRI associations were largely absent and hence are not shown on the brain surface (see Supplement Table S3 for detailed results).

First, we confirmed that the computed metastability metric captured genuine neural signal rather than statistical artifacts by comparing the empirical values against phase-randomised surrogate data that preserves spectral properties while destroying interregional phase relationships (Figure 2B). Empirical fPET and fMRI metastability values significantly exceeded phase-randomised null value (*t*(85) = 45.3, p < 0.001, Cohen’s d = 4.9 and *t*(85) = 50.1, p < 0.001, Cohen’s d = 5.8). These findings confirm that the metastability metric captures coordinated network dynamics arising from interregional coupling, rather than common statistical properties of the measured timeseries.

We compared metastability between fPET and fMRI using GLMs (Figure 2A). The two modalities exhibited significantly different network architectures across metrics (all p < 0.002). Compared to fPET, fMRI showed significantly higher metastability and global and local efficiency in absolute terms. However, the integration-segregation balance index (ISBI) was higher in fPET than fMRI, indicating that the glucodynamic signal is a closer-to-optimal balance than the haemodynamic signal. fPET showed a significantly higher number of network modules but lower modularity, indicating that fPET networks have more hub regions, although those hubs are less densely interconnected than the hubs of fMRI networks.

The partial correlation analyses between global MSE and the network metrics were significant in fPET at 16s, 32s and 48s (τ = 1, 2 and 3; Figure 1C). Specifically, global fPET MSE was positively correlated with metastability (r = 0.32 to 0.40, p-FDR < 0.05), indicating that higher metabolic signal complexity is associated with a more metastable network topology. MSE was also positively correlated with global (r = 0.27 to 0.36, p-FDR < 0.05) and local (r = 0.42 to 0.59, p-FDR < 0.05) efficiency, indicating that greater metabolic signal complexity relates to enhanced network efficiency at both global and local levels. The stronger association with local efficiency resulted in a negative correlation between MSE and the ISBI (r = -0.35 to -0.46, p-FDR < 0.05 at scales 2-3), suggesting that the local efficiency gains outweighed the global efficiency gains from higher entropy. This pattern indicates that greater metabolic complexity is associated with a network that is more efficient globally and even more efficient locally, effectively shifting the network balance toward local specialisation while maintaining global integration.

Non-linear models significantly improved fit for fPET MSE at 32s and 48s (*τ* = 2 and 3), revealing significant quadratic relationships for metastability, global and local efficiency (all p-FDR < 0.05). The pattern of association was “J-shaped”, indicating that particularly high MSE values are associated with heightened network flexibility and efficiency (Figure 2D).

In contrast to fPET, for fMRI, no significant linear nor non-linear relationships were observed between global MSE and the network metrics at any temporal scale (all p-FDR > 0.05).

Network-level analyses revealed that the association between fPET MSE and dynamic network topology was spatially specific (see Figure 2E; and Supplement 5). The strongest associations were observed in the Somatomotor (L-SOM: r = 0.42-0.52; R-SOM: r = 0.34-0.45), Salience Ventral Attention (L-SA: r = 0.39-0.50; R-SA: r = 0.25-0.40), Control (L-CON: r = 0.28-0.49; R-CON: r = 0.35-0.51), and Default (L-DEF: r = 0.40-0.62; R-DEF: r = 0.29-0.55) networks across fPET MSE windows of 16, 32 and 48s (τ = 1, 2 and 3; all p-FDR < 0.05). Conversely, the Limbic and Temporal Parietal networks showed predominantly non-significant associations. These findings indicate that metabolic complexity is most strongly tied to the dynamic topology of unimodal and higher-order association networks. For fMRI, no consistent network-level effects were observed, with only isolated effects in the Somatomotor, Visual and Temporal Parietal networks at 4s (*τ* = 4), where higher MSE was associated with lower local efficiency.

Together, these findings demonstrate that metabolic complexity and dynamic network topology are coupled, whereas for the haemodynamic signal they represent distinct, decoupled dimensions of brain function. Specifically, higher metabolic complexity across temporal scales is associated with networks that are more flexible (metastable), locally specialised and globally integrated. The divergence in results between the imaging modalities likely reflects the underlying neurovascular and metabolic processes characterised by the metabolic fluctuations of fPET compared to the haemodynamics measured in fMRI.

### MSE Associations with Cortical Thickness, Glucose Metabolism and Perfusion

We next examined whether MSE was associated with cortical thickness, baseline CMR_GLC_ measured across the scan, and CBF. After applying outlier exclusion criteria (>3 SD from the group mean), two participants were excluded from CMR_GLC_ analyses and one from CBF analyses. For the remaining participants, the cohort exhibited physiologically normal values across measures (Figure 3A). Mean cortical thickness was 2.4 mm (SD = 0.13), CMR_GLC_ 3.3 mg/100g/min (SD = 1.03), and CBF 65.6 mL/100g/min (SD = 20.3), consistent with population norms for healthy adults [55–58].

**Figure 3.**
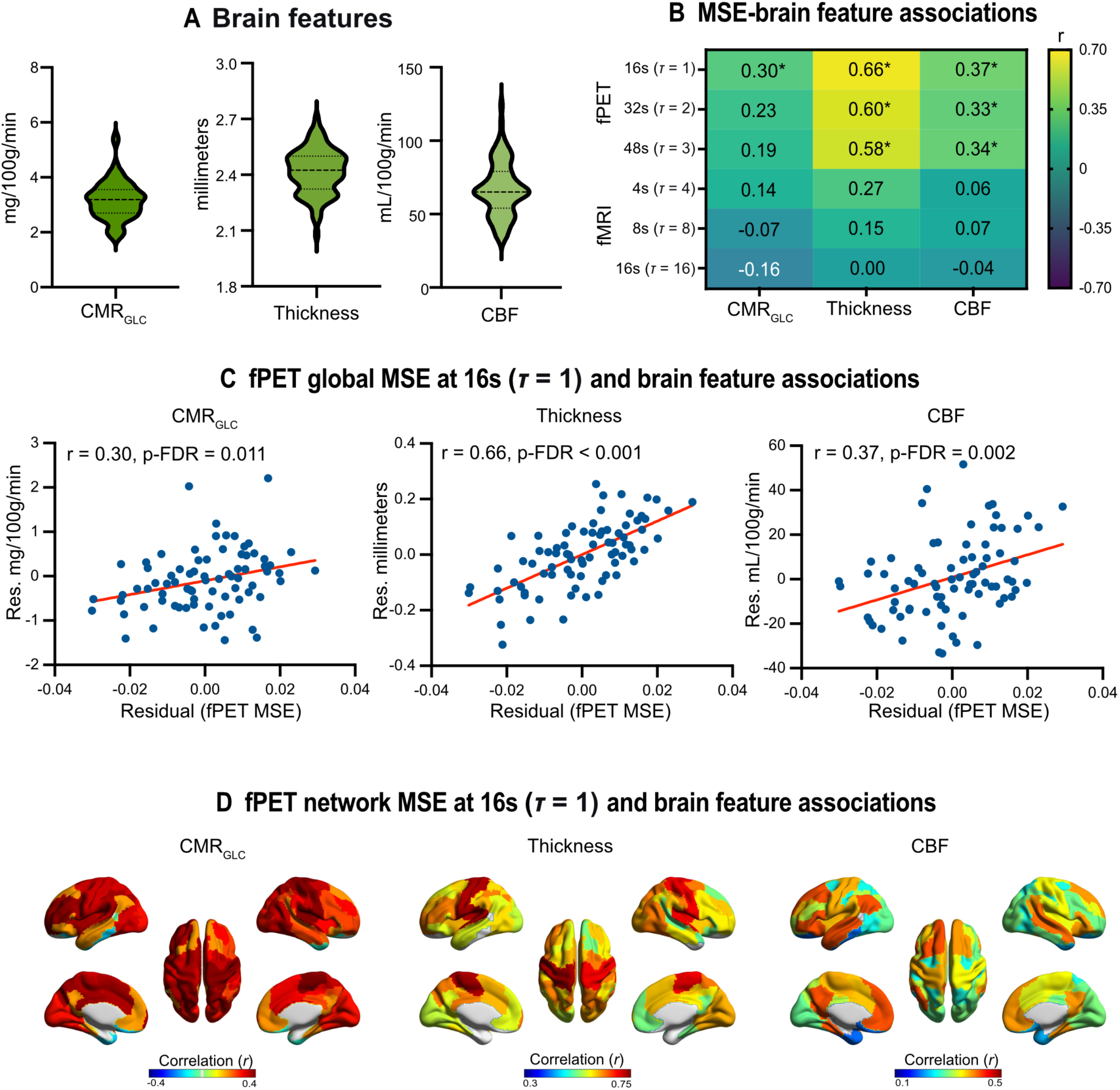
Multiscale entropy (MSE) associations with cortical thickness, cerebral metabolism and blood flow. (A) Sample distribution for whole brain cerebral metabolic rate of glucose (CMR_GLC_), cortical thickness and cerebral blood flow (CBF) and (B) their associations with MSE, controlling head motion. In (B), heatmap shows correlation coefficients (r) with asterisk (*) indicating significance at p-FDR < 0.05. Significant associations were found for fPET MSE with brain features across Scales 16s, 32s and 48s (*T* = 1, 2 and 3), but not for fMRI MSE at 4s, 8s, nor 16s (*T* = 4, 8 and 16). (C) Significant linear relationships between fPET MSE and CMR_GLC_ cortical thickness and CBF. Values are MSE residuals after controlling for motion. (D) Associations between network-level fPET MSE and CMR_GLC_ cortical thickness and CBF. fMRI associations were largely absent and hence are not shown on the brain surface (see Supplement Table S4 for detailed results). Across temporal and spatial scales, PET MSE showed positive associations with CMRglc, cortical thickness, and CBF, indicating that higher metabolic signal complexity is associated with greater glucose metabolism, thicker cortex, and higher perfusion.

To test our hypothesis that greater MSE is associated with greater cortical thickness and higher rates of metabolism (CMR_GLC_ and CBF), we undertook partial correlation analyses controlling for head motion (Figure 3B-C). For fPET, greater global MSE was associated with greater cortical thickness and CBF across all temporal windows. Greater fPET MSE was also associated with higher CMR_GLC_ and CBF at 16s (*T* = 1). In contrast, global fMRI MSE showed no significant associations with cortical thickness, CMR_GLC_ and CBF (all p-FDR > 0.05).

Network-level analyses revealed that the associations between fPET MSE and brain features were spatially specific (Figure 3D; also see Supplement 6). The strongest positive correlations were observed for MSE and cortical thickness in the Somatomotor (L-SOM: r = 0.73, R-SOM: r = 0.70; FDR p < 0.001), Visual (R-VIS: r = 0.61), and Dorsal Attention networks (L-DA: r = 0.59). In contrast, correlations in the bilateral Limbic (L-LIM: r = 0.09, R-LIM: r = 0.19) and left Temporal Parietal (r = 0.23) networks were non-significant. This pattern was consistent across fPET periods of 16-48s (*τ* = 1, 2 and 3). For fMRI, positive associations between MSE and cortical thickness were found in the bilateral Somatomotor (r = 0.35 and r = 0.31) and right Visual (r = 0.29) networks only. These results are consistent with previous fMRI complexity work indicating localised associations with cortical thickness [59, 60]. None of the fMRI MSE and CMR_GLC_ nor CBF correlations were significant.

Complementing the subject-level analyses, group-level spatial correlation analyses of independent brain maps sourced from *neuromaps* [61] revealed that MSE is associated with macroscale cortical architecture across both modalities. For fPET at 16s (*τ* = 1) and fMRI at 8s (*τ* = 8), group-averaged MSE maps showed highly significant spatial correlations with the independent sample maps of cortical thickness (fPET: r = 0.88, p-FDR < 0.05; fMRI: r = 0.86, p-FDR < 0.05) and myelin ratio (fPET: r = 0.82, p-FDR < 0.05; fMRI: r = 0.87, p-FDR < 0.05). In contrast, no significant correlations were observed for either modality with histological microstructural features, including mPC1 (r = -0.004, p-FDR > 0.05), which indexes depth-dependent laminar cell density, nor G1 (r = -0.06, p-FDR > 0.05), which captures the primary sensory-to-association hierarchy of cellular packing profiles. Similarly, group-level CMRO_2_ [62] also sourced from *neuromaps*, showed no significant correlation (both modalities, r = -0.02, p-FDR > 0.05). Notably, while simultaneously-acquired *subject*-level CMR_GLC_ and CBF were significantly associated with fPET MSE, the *group*-level CMRO_2_ map from the independent *neuromaps* sample did not survive spatial permutation testing in either modality.

Taken together, these findings suggest that MSE spatially aligns with macro-structural tissue properties (myelination and cortical thickness) rather than fine-scale microstructural histological gradients or oxygen consumption. For fMRI, haemodynamic complexity-cortical thickness associations are isolated to sensory networks. In contrast, for metabolic complexity, strong spatial correspondence with independent structural maps, combined with the subject-level global and network associations with cortical thickness, CMR_GLC_ and CBF, provides validation that signal complexity reflects biologically meaningful variation in the brain’s cortical and metabolic architecture.

### Age Differences in MSE

To test our hypothesis that MSE declines in ageing, we compared global MSE between younger (n = 39) and older (n = 45) participants using general linear models with head motion as a covariate. For fPET, we observed significant age group differences across 16, 32 and 48s windows (*τ* = 1, 2 and 3; (*F*(2, 81) = 21.6 to 35.6; all p-FDR < 0.05), with older people having lower global MSE than younger people (Figure 4A). In contrast, for fMRI, no significant age group differences were observed at any temporal scale (*F*(2, 81) = 0.6 to 1.9, all p > 0.15). These findings indicate that metabolic complexity captures age-related differences in metabolic brain dynamics, whereas haemodynamic complexity does not.

**Figure 4.**
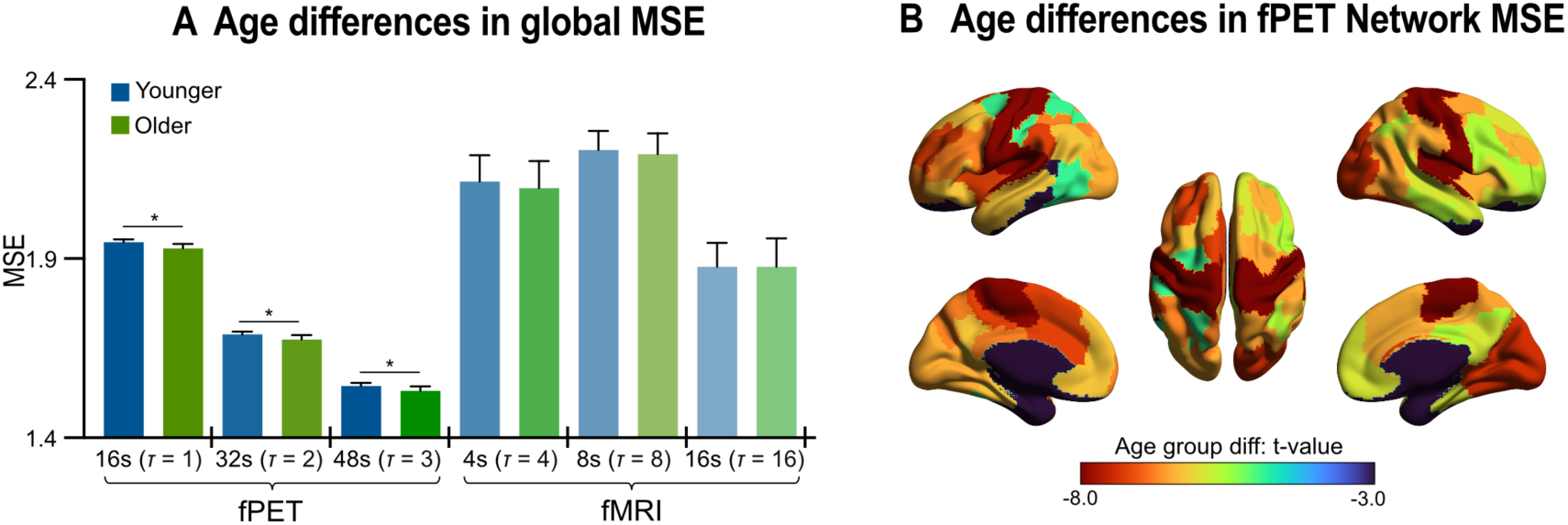
Multiscale entropy (MSE) and age. (A) Mean (SE error bars) MSE for younger and older adults. Younger adults had higher fPET global MSE than older adults at all scales (p-FDR < 0.05). (B) Age group differences in network-level fPET MSE at 16s (*T* = 1); negative *t* = younger > older. Age group differences in fMRI network-level MSE were largely absent and hence are not shown on the brain surface (see Supplement Table S5 for detailed results). Head motion was included as a covariate in the age group analyses. Asterisk (*) indicates significant effect at p-FDR < 0.05.

To assess whether this lower fPET MSE localised to specific functional systems, we conducted GLM analyses at the network level. For fPET, significant lower MSE was observed in older people, with the strongest effects in the Somatomotor, Salience Ventral Attention, Control and Default networks across all temporal scales (all p-FDR < 0.05; Figure 4B and Supplement 7). In contrast, the bilateral Limbic and left Temporal Parietal networks showed no significant age differences (p-FDR > 0.05). For fMRI, consistent with the global MSE results, no network survived FDR correction for age-related differences, with some networks showing nominally significant effects at 4s (*τ* = 4) before FDR correction (e.g., L-SOM: p = 0.004; R-SOM: p = 0.030). These findings indicate that metabolic complexity captures a spatially widespread, functionally organised metabolic ageing signature, particularly in unimodal and higher-order association networks, that is absent in haemodynamic complexity.

### MSE and Cognition

Lastly, we used correlation analysis to test our hypothesis that greater MSE is associated with better cognitive performance. Cognitive performance was assessed using four standardised tasks measuring domains validated in ageing research (see Methods): verbal memory (delayed recall from the Hopkins Verbal Learning Test), cognitive control (reaction time during task-switching), response inhibition (reaction time during a stop-signal task), and visuospatial processing speed (number of correct responses during digit substitution). For the correlation analyses, the reaction time measures were multiplied by -1 so that higher scores correspond to better (faster) performance.

The sample distributions of the cognitive measures are shown in Figure 5A. As expected for a community-dwelling, cognitively healthy cohort, the distributions were approximately normal, with no indication of floor or ceiling effects. For fPET, significant positive correlations were observed between MSE and cognitive domains across all temporal scales (Figure 5B, left). Specifically, fPET MSE at 16s (*τ* = 1) was positively correlated with episodic memory (r = 0.34), task switching reaction time (r = 0.35) and processing speed (r = 0.57), all p-FDR < 0.05. These associations remained significant at 32 and 48s (*τ* = 2 and 3). In contrast, no significant correlations were observed between fMRI MSE and any cognitive measure (Figure 5B, right), and no non-linear (quadratic) relationships were observed for either modality. To determine whether these associations were independent of age, we repeated the analyses with age included as a covariate. Controlling for age attenuated all previously significant fPET MSE-cognition correlations (all non-significant after FDR correction), suggesting that metabolic complexity is associated with age-related changes in brain dynamics rather than isolated cognition effects per se.

**Figure 5.**
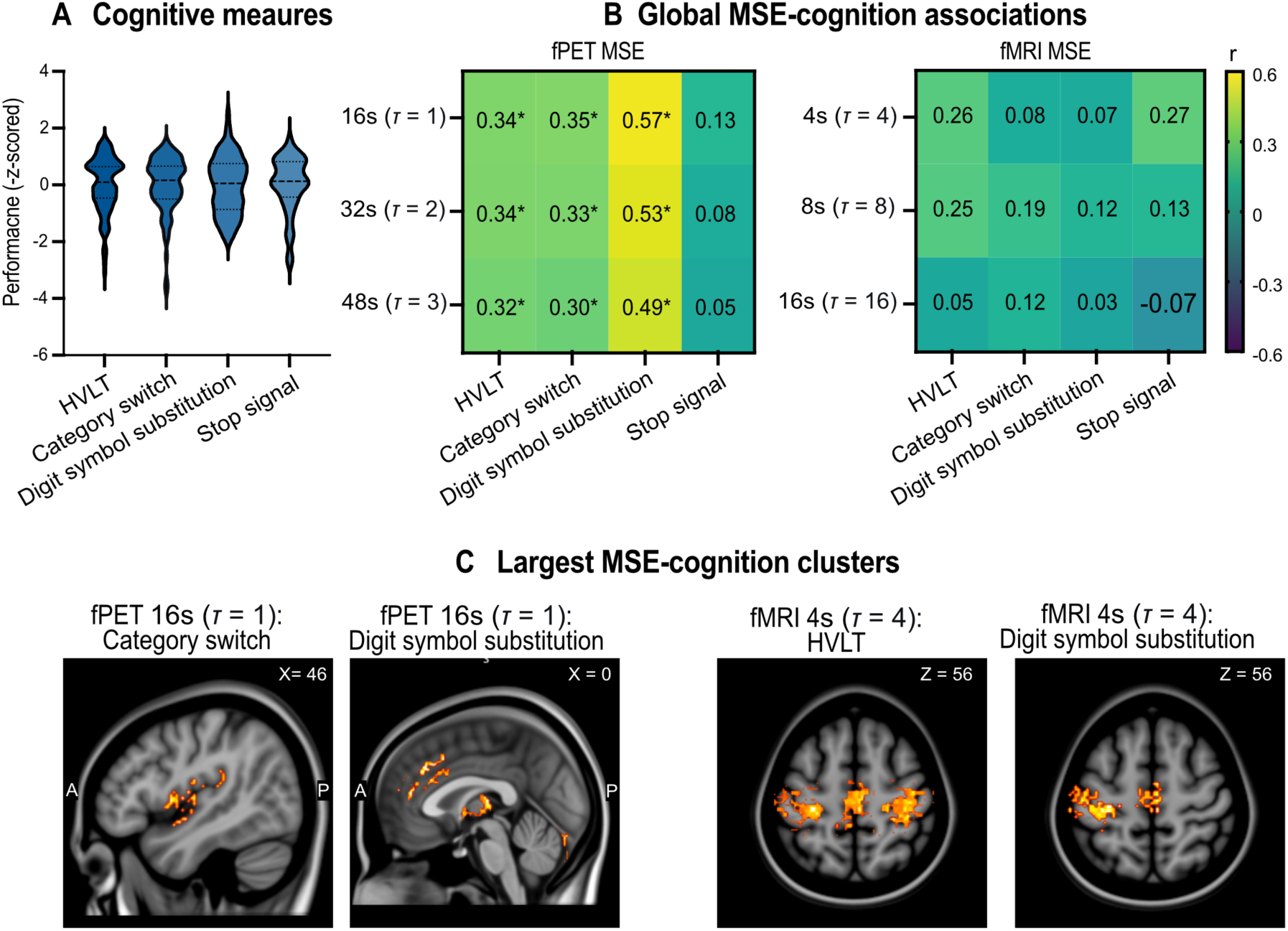
Multiscale entropy (MSE) and cognition. (A) Sample distribution of cognitive measures: HVLT delayed recall, category switching RT, digit symbol substitution, and stop signal RT. (B) Correlation analyses between global MSE and cognitive test performance. Heatmaps show correlation coefficients (r) for fPET MSE (left) and fMRI MSE (right). Significant associations were found for fPET MSE across Scales 1-3 but not for fMRI MSE at any scale. (C) Largest significant clusters identified from voxel-level MSE-cognition analyses. Significant clusters were found in both modalities (fPET: 71 clusters, ranging from 32-552 voxels; fMRI: 17 clusters, ranging from 35-4,991 voxels), with the largest fPET clusters localised to the thalamus for digit symbol substitution (552 voxels) and the right central opercular cortex for category switching (255 voxels); and for fMRI, the largest cluster was in the precentral gyrus for HVLT (4,991 voxels), with additional clusters in the postcentral gyrus for both category switching (627 voxels) and digit symbol substitution (997 voxels). All associations were no longer significant when age was included in the models, with the exception of four fMRI clusters (35 to 1,039 voxels), indicating that MSE-cognition relationships are largely explained by age-related variance. Head motion was included as a covariate in cognition analyses. Asterisk (*) indicates significant effect at p-FDR < 0.05.

Because previous fMRI and EEG work in MSE and cognition has used voxel-level clustering or SPM-based analysis (e.g., [35, 63]), we reasoned that the lack of association between MSE and cognition in our fMRI data may reflect the spatial scale of our analysis. Hence, as a final step, we investigated MSE-cognition associations at the voxel-level. We conducted voxel-wise GLM analyses between MSE and cognition with permutation-based cluster extent correction. An initial voxel-wise cluster identification threshold of p < 0.005 was applied, with permutation testing (1,000 permutations) used to determine cluster extent thresholds (α = 0.05).

Significant clusters were found in both modalities. For fPET, 73 clusters, ranging from 32 to 552 voxels, were found for the 16, 32 and 48s periods (*τ* = 1, 2 and 3), predominantly for digit symbol substitution (55 clusters: 14 at 16s, 20 at 32s and 21 at 48s) and category switching (16 clusters: 3 at 16s; 6 at 32s and 7 at 48s). HVLT delayed recall (32 voxels) and stop signal (36 voxels) yielded a single cluster at 48s and 32s, respectively (see Supplement 8). The clusters were spatially distributed across a diverse range of cortical and subcortical regions, encompassing the thalamus, insula, central opercular cortex, cingulate cortex, paracingulate gyrus, temporal pole, prefrontal cortex, parietal lobules, and cerebellum. The largest clusters were all found at 16s (*τ* = 1) and localised to the left thalamus for digit symbol substitution (552 voxels) and the right central opercular cortex for category switching (255 voxels) (Figure 5C). Additional fPET clusters included the left insula for digit symbol substitution (182 voxels), the right paracingulate gyrus for digit symbol substitution (219 voxels), and the anterior cingulate cortex for category switching (146 voxels).

In contrast, fewer but mostly larger and spatially focussed clusters were found for fMRI, with 17 clusters ranging in size from 35 to 4,991 voxels. Importantly, all fMRI clusters were localised exclusively to the pre- and post-central gyri at 4s and 8s (*τ* = 4 and 8; see Supplement 7 and 8). The largest fMRI clusters were found at 4s in the right precentral gyrus for HVLT (4,991 voxels), the left postcentral gyrus for digit symbol substitution (997 voxels) and the right postcentral gyrus for category switching (627 voxels).

When the voxel-level analyses were repeated with age as a covariate, all fPET clusters identified without age were no longer significant, indicating that voxel-level fPET MSE-cognition relationships are driven by age-related variance (see Supplement 8). For fMRI, 13 of 17 clusters (76%) were eliminated when controlling for age, with the number of voxels reducing by 79% across the clusters (see Supplement 9). Four clusters survived age correction: HVLT at 4s (2 clusters; 1,039 and 454 voxels), category switching at 8s (1 cluster; 35 voxels), and stop signal at 4s (1 cluster; 229 voxels). The surviving fMRI clusters were localised to the right precentral gyrus (3 clusters) and left postcentral gyrus (1 cluster).

Taken together, these results suggest that brain signal complexity associations with cognitive performance are largely accounted for by age variance across temporal and spatial scales. Notably, while metabolic complexity showed robust relationships across spatial scales, haemodynamic complexity effects were only detected at the voxel level. Furthermore, the spatially restricted nature of fMRI clusters in pre and postcentral gyri contrasted with the widespread cortical and subcortical distribution of smaller fPET clusters, suggests that metabolic and haemodynamic signal complexity capture fundamentally different topologies of cognitive ageing relationships.

### Sensitivity analyses

We undertook sensitivity analyses using alternative embedding and entropy measures, including sample entropy, Shannon entropy, Brain Entropy (BEN), and Fuzzy entropy. A description of how these measures each capture different aspects of brain signal complexity, together with the detailed results of the sensitivity analyses, are available in Supplement 10. Briefly, all fPET and fMRI entropy measures were less irregular than phase-randomised surrogates. All fPET entropy measures showed positive associations with metastability, efficiency, cortical thickness, CMR_GLC_, CBF and cognition, as well as significantly lower values in older than younger people. By comparison, associations were largely absent for all fMRI entropy measures. Only Shannon entropy of the fMRI signal showed age differences, with higher values in older people. Overall, the sensitivity analyses support our main findings of modality differences in MSE and their associations with network metastability, ageing and cognition.

## Discussion

Here, we demonstrate that the brain’s metabolic system exhibits temporal complexity consistent with the theories of coordination dynamics [2, 12] and metastability [1, 2, 5]. Our primary findings were four-fold. First, we identified entropy signal in the temporal dynamics of the glucose (fPET) timeseries, thereby confirming that the brain’s glucose energetic framework exhibits temporal complexity. Second, metabolic complexity mapped to network topology. Across unimodal and higher-order association networks, greater glucodynamic complexity was associated with more flexible network dynamics (metastability), greater cortical structure (thickness, myelin) and higher rates of metabolism (CMR_GLC_, CBF). Third, glucodynamic complexity was sensitive to phenotype, showing robust lower global and network values in older compared to younger adults. Although greater global and network-level glucodynamic entropy was associated with better cognitive performance, and local clusters of both fPET and fMRI entropy were associated with cognition, those associations were almost entirely accounted for by age. This indicates that glucodynamic and haemodynamic complexity track age-related brain changes but are not separately related to cognitive performance. Fourth, haemodynamic complexity was unrelated to age, metabolism and network dynamics. Taken together, these results build on a growing body of work indicating that the dynamics of cerebral glucose metabolism are critical to the organisation and function of the brain [20, 27–31, 64, 65]. They also suggest that the time-variant complexity of glucose metabolism can complement standard linear and static measures of brain activity and may be suited as a biomarker of brain ageing.

Our findings reframe the role of cerebral glucose metabolism in brain organisation and function. Rather than acting as a passive energetic pool awaiting neural demand, or a simple downstream consequence of neural firing, the glucodynamic signal carries organised temporal structure and is tied to the network-level flexibility that underlies adaptive brain function. In other words, metabolism is an *active* contributor to the brain’s coordination dynamics, not simply the metabolic substrate that supports them. This interpretation has several conceptual implications. Firstly, it removes any hierarchy in which metabolism is subordinate to neural activity. Secondly, our data show that glucodynamic complexity is age-sensitive in a way that haemodynamic complexity is not. These results are difficult to reconcile with a purely downstream energetic support role in neural activity. Rather, they suggests that the metabolic and haemodynamic systems are not interchangeable measures of the same underlying processes. Thirdly, our results also imply that energetic and information-processing models of brain function, often treated as separate explanatory frameworks [66], are in fact two perspectives on the same dynamic system [17]. The brain does not simply process information using energy; it processes information through the dynamic organisation of its energy use. Under this view, measures like entropy of metabolic signals are not purely neurophysiological markers but direct measures of the temporal structure of brain metabolism, as would be expected if metabolism is a constituent of, rather than a support for, brain network coordination dynamics.

Across temporal scales, greater metabolic signal complexity was associated with greater network metastability and local and global efficiency, with the strongest effect obtained for local efficiency. This pattern is consistent with the view that greater regional complexity enables dynamic network configurations, reflecting information-rich signals that support specialised local processing and efficient global integration [5, 41]. The relatively greater increase in local efficiency with higher entropy may reflect enhanced specialised processing, while the absolute increase in global efficiency ensures information can still be integrated across networks when needed [13, 15]. Theoretically, metastability reflects the brain’s capacity to flexibly transition between different network configurations without becoming trapped in a single state [1–3, 5, 9]. Our results align with prior fMRI [8, 40, 41] and EEG [67] work, suggesting that higher glucodynamic entropy supports this metastable regime, allowing for the rapid state switching and global integration that are hallmarks of healthy brain function. Moreover, the consistency in results across metabolic entropy windows appears to reflect an optimised network configuration where the brain can utilise dynamic glucose signals to efficiently process information at multiple temporal scales simultaneously [68].

We also found that greater glucodynamic complexity is closely associated with greater cortical thickness and myelin, suggesting that metabolic complexity captures the information-processing capacity represented by available cortical tissue. Our results also suggest that higher baseline glucose metabolism is associated with more complex, information-rich neural dynamics. These results complement a recent report showing that, when glucose drops below a critical threshold (i.e., hypoglycaemia), the brain’s ability to maintain complex electrophysiological dynamics is impaired [69]. They are also consistent with the concept that higher signal complexity reflects more efficient CMR_GLC_ and CBF regulation [70]. Together, these results establish brain metabolic complexity as a neurobiologically meaningful metric that integrates structural, metabolic and network features of the brain.

A strong and consistent finding of this study was the robust lower glucodynamic complexity in older people. Entropy of electrophysiological and haemodynamic signals has previously been linked to healthy ageing [35, 46, 71, 72]. The current findings extend this to brain metabolism by demonstrating that the brain’s glucodynamics lose complexity in older people. We speculate that this loss of energetic complexity contributes to the “brain energy crisis” in older adults, in which a progressive deficit occurs in the ability to generate sufficient ATP to meet metabolic demands [73]. It is driven by an impairment in neural glucose uptake and utilisation [73], compromising synaptic transmission and network connectivity, thereby setting the individual on a trajectory towards age-related decline and neurodegeneration. The loss of energetic complexity in older people observed here likely occurs concomitantly with age-related reductions in brain structure and function, including loss of neural plasticity, reduced neurovascular coupling, impaired insulin sensitivity, and reductions in metabolic active tissue from grey matter atrophy [36, 41, 44, 74, 75]. Given our finding that glucodynamic complexity is linked to neuro-morphological and metabolic features, this raises the question of whether the reduction of glucodynamic complexity leads to, or is caused by, these underlying biological changes. Future studies should explore this question.

In contrast to metabolic complexity, haemodynamic complexity demonstrated minimal differences between younger and older adults. While these results initially appear inconsistent with prior reports of age-related declines in fMRI and EEG entropy [35, 36, 44, 46, 76], they become interpretable when considering the non-linear physiological properties of the BOLD signal. Surrogate testing demonstrated that empirical fMRI entropy fell significantly below phase-randomised surrogates. Because phase-randomised surrogates preserve the linear autocorrelation and spectral profile of the BOLD signal, thereby fully accounting for linear haemodynamic response function (HRF) smoothing, empirical values falling below these surrogates reflect genuine non-linear neurovascular phase coupling and deterministic state dynamics [54]. This non-linear neurovascular constraint may render BOLD complexity less sensitive to normative ageing or structural variation than metabolic measures. Furthermore, BOLD fMRI does not index neural activity directly; instead, it reflects a composite neurovascular response driven by baseline venous blood volume, cerebral blood flow, and oxygenation, each influenced by intrinsic vascular dynamic processes, such as vasomotion [77, 78]. Finally, methodological heterogeneity across the literature likely accounts for divergent findings: a recent review of 92 fMRI entropy studies [37] highlighted that only a small subset constructed surrogate null distributions, with substantial variation in entropy algorithms, spatiotemporal scales, and acquisition protocols. Additionally, our extended 20-minute resting-state fMRI acquisition provides significantly higher estimation stability for multiscale entropy than the shorter or task-based scans predominantly featured in prior work [37].

A central question of this study was whether metabolic complexity predicts cognitive performance. We found that the signal complexity-cognition associations were largely explained by age variance. While it may be the case that glucodynamic variability is unrelated to cognitive performance independently of age, it is also possible our protocol may have been insensitive to cognition-related temporal complexity. In particular, the 16s temporal resolution of the fPET may have been inadequate to detect temporal complexity at the scale relevant for cognitive performance. There are some hints in the data supporting this: prior to correction for age, significant fPET MSE-cognition relationships were obtained for memory and proactive control (category switch) but not reactive control (stop-signal reaction time). While all four domains rely on slow rhythms (e.g., theta, alpha), reactive control additionally requires fast-wave (beta) activity [79, 80]. Changes in glucodynamic activity occur within milliseconds after neuronal firing [81], at a rate too fast to detect with our sampling rate. While detecting such fast glucodynamics with FDG-PET is likely outside the realm of possibility, it is currently possible to detect faster dynamics with more sensitive next-generation PET scanners, that have temporal resolutions in the order of 1-4sec [82]. Future fPET studies that use improved temporal dynamics may be able to reveal stronger cognitive associations.

Several limitations should be noted. First, as mentioned above, the fPET data were acquired with a temporal resolution of 16s per frame, which is markedly slower than the fMRI. This coarse temporal sampling restricts our ability to capture fast, high-frequency neural dynamics reflected in the metabolic domain. It also restricted our ability to compare multiscale entropy between imaging modalities on a range of consistent scales. Future studies should explore glucodynamic multiscale entropy at faster temporal scales. For example, it will be important to test if metabolic complexity exhibits a similar inverted-U shape across scales that is seen in the haemodynamic signal.

In conclusion, this study demonstrates that the complexity of cerebral metabolism is tied to the brain’s synchronised dynamics. Multiscale entropy of the metabolic signals reflects the brain’s capacity to flexibly transition between network configurations without becoming locked into a single state, consistent with the theories of coordination dynamics and metastability. Entropy of dynamic glucose use also maps to the brain’s structural and metabolic architecture, underscoring its neurobiological basis, and is a highly sensitive marker of brain ageing. In contrast, haemodynamic multiscale complexity was largely unrelated to network dynamics and insensitive to age. These results add to a growing literature indicating the temporal dynamics of metabolic signals carry important biological and functional information that is complementary to the information contained in other imaging modalities. We have shown that the brain’s adaptive capacity is tied to the complexity of its neural activity, and that multiscale metabolic entropy relates to brain network organisation, brain ageing and cognitive decline. The complexity of glucodynamic signals and dynamic metabolic networks offer a unique way to study the whole brain and offer promise as a biomarker in ageing and disease.

## Methods

### Ethical Considerations

The Monash *MetConn* simultaneous PET/MR dataset was used for the study [20, 28, 83]. Primary data collection and analysis were conducted in accordance with the Australian National Statement on Ethical Conduct in Human Research and the 1964 Declaration of Helsinki. Institutional ethical approval was provided by the Monash University Human Research Ethics Committee (Project ID: 26415). Monash Health Principal Medical Physicist approved the use of ionizing radiation based on the Australian Radiation Protection and Nuclear Safety Agency Code of Practice (2005). Participants provided informed consent to participate.

### Participants

Local advertising was used to recruit 90 participants from the general community. Participants were screened for the capacity to provide informed consent. Exclusion criteria included neurological or psychiatric illness, a diagnosis of diabetes, or dementia. Women were screened for current or suspected pregnancy. Participants received a $100 voucher for completing the study. Six participants were excluded from further analyses due to head motion (n=2), incomplete PET scan (n=2) or being > 3 SDs in MSE analyses (N = 2). The final sample included 84 participants (see Supplement 1 for demographics).

### Demographic and Cognitive Variables

Prior to the scan, participants completed a demographic questionnaire and a battery of cognitive tests, comprised of domains validated in research on ageing [84] (see Supplement 2 for details): delayed recall from the Hopkins Verbal Learning Test [85]; reaction time in a task switching test to index cognitive control [86]; reaction time in a stop signal task to measure response inhibition [87]; and the number of correct responses in a digit substitution task to measure visuospatial performance [88].

### MR/PET Data Acquisition

Participants underwent a 90-minute simultaneous MR/PET scan in a Siemens Biograph 3-Tesla molecular MR scanner. They were cannulated in the vein in each forearm. Half of the 260 MBq FDG tracer was administered via the left forearm as a bolus at the start of the scan. The remaining 130 MBq was infused over 50 minutes at 36ml/hour. This infusion protocol maximises the signal-to-noise ratio over the scan [89].

Participants were positioned supine in the scanner bore with their head in a 32-channel radiofrequency coil. The scan sequence started with non-functional MRI scans during the first 12 minutes, including a T1 3DMPRAGE (TA = 3.49 min, TR = 1,640ms, TE = 234ms, flip angle = 8°, field of view = 256 × 256 mm^2^, voxel size = 1.0 × 1.0 × 1.0 mm^3^, 176 slices, sagittal acquisition) and T2 FLAIR (TA = 5.52 min, TR = 5,000ms, TE = 396ms, field of view = 250 × 250 mm^2^, voxel size = .5 × .5 × 1 mm^3^, 160 slices). List-mode PET (voxel size = 2.3 x 2.3 x 5.0mm^3^) sequences commenced at the start of the scan and T2* EPI BOLD-EPI sequence at 13 minutes (TA = 40 minutes; TR = 1000ms, TE = 39ms, FOV = 210 mm^2^, 2.4 × 2.4 × 2.4 mm^3^ voxels, 64 slices, ascending axial acquisition), acquired while participants watched a movie of a drone flying over the Hawaii Islands. At 53 minutes, a 5-delay pseudo-continuous arterial spin labelling (pcASL) scan was initiated (TR = 4,220 ms; TE = 45.46 ms; FOV = 240 mm; slice thickness = 3 mm; voxel size 2.5 x 2.5 x 3.0 mm^3^). Post labelling delays were 0.5, 1.0, 1.5, 2.0, and 2.5s and duration of the labelling pulse was 1.51s.

Plasma radioactivity levels were measured during the scan from 5ml blood samples taken from the right forearm, every 10-minutes, for a total of nine samples. The samples were spun in a centrifuge at 2,000 rpm for 5 minutes. 1,000-μL of plasma was placed in a well counter for four minutes, with the start time, number of counts, and counts per minute recorded.

#### PET and fMRI Processing and Brain Feature Extraction

The pial and white matter surface of the T1 images were quality checked and corrected where needed and the images registered to MNI152 space . For the BOLD-fMRI data, T2* images underwent brain extracted, unwarping and motion correction. Motion was corrected using six rotation and translation parameters [90]. The images were also temporally detrended, normalised to MNI space and smoothed at 8mm FWHM [90]. To quantify head motion, framewise displacement was calculated for each participant and used as a covariate in the primary analyses.

The list-mode PET data were binned into ∼344 3D sinogram frames of 16s intervals, corrected for attenuation and reconstructed using Ordinary Poisson-Ordered Subset Expectation Maximization algorithm (3 iterations, 21 subsets) with point spread function correction. The reconstructed DICOM slices were converted to NIFTI format with size 344 × 344 × 127 (size: 1.39 × 1.39 × 2.03 mm^3^) for each volume. A single 4D NIFTI volume was created by concatenating the 3D volumes, which were motion [90] and partial volume corrected [91], applying a 25% grey matter threshold [92]. The PET images were also spatially smoothed [93] using a Gaussian kernel with a full width at half maximum of 8mm and co-registered to MNI space.

The *CONN* toolbox [94] was used to denoise the fPET and fMRI data of each participant by removing white matter and CSF confounds. A bandpass frequency filtered of 0.01 Hz to 0.1 Hz was used for the fMRI data. As grey matter uptake of FDG is relatively uniform across the brain, regions are likely to show some degree of temporal complexity, particularly if the initial steep rise of the tracer uptake is included in the timeseries. To minimise this issue, the first 10 minutes of the timeseries was excluded to remove the initial uptake and to reflect the time corresponding to the onset of a stable signal [89]. The resulting fPET and fMRI timeseries represent the relative FDG and BOLD signal changes across frames and volumes, respectively, from which MSE and network metrics were calculated.

### Multiscale Entropy Calculation

MSE analysis was performed on the 4D fPET and fMRI images using a coarse-graining approach [33] with the following parameters: embedding dimension (m) = 2; tolerance threshold (r) = 0.15 × standard deviation of each voxel’s time series. Scale factors (*τ*) = 1, 2, 3 and 4 for fPET and 1, 2, 3, 4, 8, 16, 32 and 48 for fMRI. To balance computational demand with statistical requirements, we computed fMRI entropy from frame 600 (matching the 10-minute mark of the fPET) for 1,200 volumes (i.e., 20 minutes).

For each scale factor (τ), the time series were coarse-grained by averaging consecutive data points in non-overlapping windows of length τ:

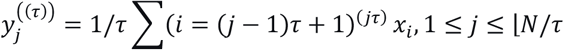

Sample entropy was then calculated for each voxel’s coarse-grained time series. For a time series of length N, vectors of length m are constructed as:

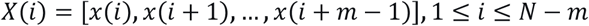

The number of vector pairs with Chebyshev distance ≤ r was counted for both m-dimensional (B) and m+1-dimensional (A) vectors, and sample entropy was calculated as:

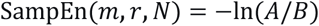

When A or B equalled 0, the entropy value was set to NaN. This procedure was repeated for all scale, resulting in 3D MSE maps per participant, each representing the temporal complexity at a different time scale. MSE values were also extracted at the regional level by computing voxel-wise mean entropy profiles across each of the 100 cortical parcels in the Schaefer atlas. Regional estimates were subsequently aggregated into 16 canonical functional networks using volume-weighted averaging, with global MSE defined as the mean entropy across these network parcels.

### Network Topology

To characterise dynamic network organisation, we computed a comprehensive set of network metrics using both the fPET and fMRI timeseries. First, regional timeseries were extracted for the 100 Schaefer regions [47].

#### Metastability

For each subject and modality, the instantaneous phase of the regional timeseries was computed using the Hilbert transform, and the Kuramoto order parameter was used to quantify global synchrony over time [9, 95]. Metastability was defined as the standard deviation of this synchrony time series. To validate that the resulting metastability values reflect genuine neural coordination rather than statistical artifacts, we performed a surrogate data analysis. For each subject and modality, 100 phase-randomised surrogates were generated that preserved the spectral power of the original timeseries while destroying interregional phase relationships. Empirical metastability values were compared against surrogate null distributions using paired t-tests, and effect sizes were quantified using Cohen’s d.

#### Modularity and Integration-Segregation

Functional connectivity matrices were constructed by computing Pearson correlations between all pairs of regional timeseries, and negative correlations were removed. Matrices were thresholded to retain only the top 15% of edges, which represents the optimal cost-efficiency point. Using the Brain Connectivity Toolbox (BCT), modularity (Q) and the number of modules were calculated using the Louvain algorithm. Additionally, global efficiency, local efficiency and the integration-segregation balance index (ISBI, global divided by local efficiency [13]) were computed from the thresholded matrices. The ISBI provides a single metric capturing the balance between global integration and local segregation. Values near 0.5 indicate optimal network configuration, while deviations suggest a bias toward either integration (ISBI > 0.5) or segregation (ISBI < 0.5).

### Brain feature extraction

#### Metabolic rate of glucose

Calculations of CMR_GLC_ were undertaken using the FDG time activity curves for the Schaefer atlas 100 regions. The FDG in the plasma samples was decay-corrected for the time between sampling and counting as the input function to Patlak models [96].

#### Cortical Thickness

Cortical thickness was extracted for the 100 regions of the Schaefer parcellation [47]. We used FreeSurfer’s *recon-all* pipeline to generate cortical surface reconstructions and vertex-wise cortical thickness maps [91]. Cortical thickness was averaged across all surface vertices belonging to each of the 100 regions.

#### Cerebral Blood Flow

To quantify cerebral blood flow, whole−brain CBF maps were calculated from perfusion weighted pcASL images [97]. Image processing included motion, distortion and partial volume correction, a macro vascular component, adaptive spatial regularisation of perfusion and T1 uncertainty. The CBF images were aligned to the anatomical T1 images and normalised to MNI152 space and values calculated for the Schaefer 100 regions for each participant.

#### Independent Sample Analysis

Group-level spatial correlations were also performed against HCP myelin, thickness and CMRO_2_ maps using the *neuromaps* toolbox [61].

### Statistical Analysis

#### MSE Surrogate Testing

To validate that MSE captured genuine temporal structure rather than spectral artefacts, we generated 100 phase-randomised surrogates for each subject, modality, and scale. Surrogates preserved the power spectrum while randomising phase, and were generated from the same coarse-grained timeseries used in the empirical analysis (see Supplement 3 for illustration). For each subject and scale, we sampled 1,000 grey matter voxels and compared the empirical regional-mean MSE against its surrogate distribution via a z-score (empirical − surrogate mean) / surrogate SD; two-tailed p-values were computed parametrically and non-parametrically. Group-level inference used one-sample t-tests on per-subject z-scores across subjects. Effect sizes were quantified as Cohen’s d on paired differences and as the group mean z. The proportion of subjects with |z| > 1.96 was recorded as a measure of consistency. Analyses were performed independently for fPET at 16-64s (*T* = 1-4) and fMRI at 1-48s (*T* = 1, 2, 3, 4, 8, 16, 32, 48).

#### MSE, Network Topology and Brain Features

For each modality and scale, we tested both linear and non-linear (quadratic) relationships between global MSE and each network metric (metastability, modularity, ISBI, local and global efficiency), and brain features (CMR_GLC_, CBF, cortical thickness) using partial correlation analysis with head motion as a covariate. Statistical significance was assessed using FDR correction across all modality-scale-metric combinations, with significance tested at p-FDR < 0.05.

#### MSE and Mapping of Independent Sample Brain Features

To validate the spatial biological basis of our MSE maps, we performed group-level spatial correlation analyses using the *neuromaps* toolbox [61]. Individual-subject fPET (16s; *T* = 1) and fMRI (8s *T* = 8) MSE maps were transformed from MNI152 space to the fsaverage 10k surface space using standard resampling algorithms. These maps were then spatially correlated against a set of established brain atlases, including cortical thickness and myelin ratio from the Human Connectome Project (HCP; [98]), the BigBrain microstructural profile and gradient maps (mPC1, G1; [99]), and cerebral metabolic rate of oxygen (CMRO_2_) map [62]. To control for spatial autocorrelation, statistical significance was assessed using spin-based permutation testing [100] with 100 spin permutations. P-values were corrected for multiple comparisons at FDR < 0.05 across all density maps.

### MSE and Cognition Associations

MSE-cognition associations were tested at the whole-brain and network levels. For each modality and scale, we tested both linear and non-linear (quadratic) relationships between global MSE and cognition (HVLT delayed recall, category switching reaction time, digit symbol substitution, and stop signal reaction time), controlling for head motion and correcting for multiple comparisons at p-FDR < 0.05.

Additionally, previous work has indicated that fMRI entropy-cognition associations can be locally driven [63]. We therefore employed a data-driven voxel-wise approach to identify MSE clusters of significant associations with cognition. We performed voxel-wise general linear model (GLM) analyses using permutation-based cluster extent correction separately for each cognitive variable. An initial voxel-wise threshold of p < 0.005 (uncorrected; t = 2.887, df ≈ 80) was applied to identify candidate clusters. Cluster extent thresholds were determined using Freedman-Lane permutation testing (1,000 permutations) to control the family-wise error rate at α = 0.05. At each permutation, we computed the maximum cluster size under the null hypothesis by permuting residuals from the reduced model (motion only) and refitting the full model (motion and cognitive variable). The 95th percentile of the resulting null distribution of maximum cluster sizes served as the critical cluster extent threshold, with only clusters exceeding this threshold considered significant. To assess the influence of age on the observed brain-cognition relationships, we repeated the analyses with age included as an additional covariate.

## Data, Code, and Materials Availability

All data and code needed to evaluate and reproduce the results in the paper are present in the paper and/or the Supplementary Materials.

## Use of AI

During the preparation of this work, the authors used Gemini to debug code and improve text. The authors reviewed and edited the output as needed and take full responsibility for the content of the published article.

## Acknowledgements

We thank Robert Di Paolo, Gerard Murray, M. Navyaan Siddiqui, Katharina Voigt, Richard McIntyre, Lauren Hudswell for their contributions to data acquisition and image reconstruction.

We also acknowledge the facilities and scientific and technical assistance of the National Imaging Facility (NIF), a National Collaborative Research Infrastructure Strategy (NCRIS) capability at Monash Biomedical Imaging (MBI), a Technology Research Platform at Monash University.

## 1. Participants – demographics and cognition

The characteristics of the whole sample (N =84), as well as the younger (N = 39) and older (N = 45) participants, are shown in Table S1. The mean age of the whole sample was 53.6 years (SD = 24.6). The proportion of women was 52%. The average years of education was 16.9. Average BMI was 25.0 kg/m^2^, resting heart rate was 78.4 BPM and systolic and diastolic blood pressure were 135.6 and 81.5 mmHg, respectively. The mean fasting blood glucose was 5.0 mmol/L.

The mean age of the younger group was 28.1 years and the older group 75.7 years. The proportion of women in the younger group (54%) and older group (51%) was not significantly different. The average years of education was higher in the younger (18.1) than the older group (16.4).

The older group had significantly higher mean systolic blood pressure than the younger group (149.87mmHg vs 119.18mmHg). The older group also had a higher fasting blood glucose level (5.18 vs 4.78 mmol/L). The older group also has significantly worse performance on the HVLT (7.36 vs 9.28), category switch reaction time (2.15 vs 1.44s), stop signal RT (0.60 vs 0.54s), and digit symbol substitution (27.7 vs 62.7) performance.

**Table S1.**
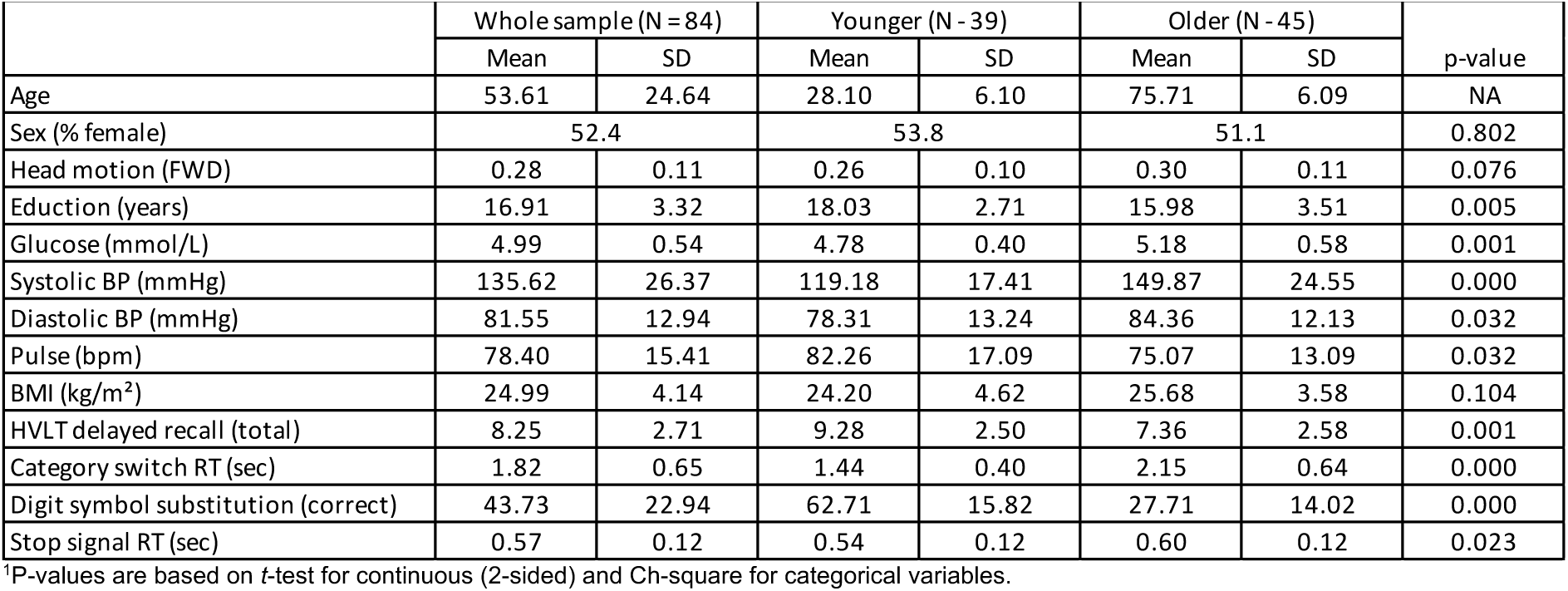
Demographics for the whole sample and comparison of older and younger groups. Continuous variables are mean (standard deviation); categorical variables are percentage.

## 2. Cognitive tests

### Hopkins Verbal Learning Test (HVLT)

The HVLT is a three-trial list learning and free recall task. The learning trials comprised 12 words, four words from each of three semantic categories. Approximately 20–25 minutes after the learning trials, participants completed delayed recall and recognition trials. The delayed recall required free recall of any of the 12 words. The recognition trial comprised 24 words, including the 12 target words and 12 false-positives, six semantically related, and six semantically unrelated. Delayed recall was calculated as the total words recalled.

### Task Switching

For task switching, a computerised test was used in which participants were presented with a word and had to perform a categorisation task. The categorisation task was dependant on two cues that appeared on screen across the trials. One cue was a heart symbol, for which participants were asked to categorise the word presented via a key press as either a LIVING or a NON-LIVING object. If the cue was an arrow-cross, participants were asked to categorise the word as either BIGGER or SMALLER than a basketball. The cue was randomised across trials. Half the trials were switch trials and half were non-switch trials. Half the switch and non-switch trials was congruent in the key presses for either task, half was incongruent. The task switching measure was latency of correctly responding to a switch trial.

### Stop Signal

The stop signal trial was a computer-based test in which participants were required to press the left response key if an arrow on screen pointed left and the right response key if the arrow pointed right. If a signal beep sounded, participants were instructed to stop their response. The delay between presentation of an arrow and signal beep started at 250ms and was altered up or down by 50ms based on performance. The delay increased up to 1150ms if the previous stop signal trial was successful and decreased to 50ms if the previous trial was unsuccessful. The stimulus onset asynchrony between the onset of a fixation circle at the start of each trial was 2000ms. Reaction time in the stop signal trials was recorded.

### Digit Symbol Substitution

A computer-based task presenting participants with a matrix of 18 column and 16 rows. Participants were required to translate symbols shown above the matrix in a key into digits in the matrix. The trial lasted two minutes. Performance was measured as total count of correct responses.

### Digit Span

A measure of verbal short term and working memory used in two formats: Forward and backward digit span. Participants were presented with a series of digits and are asked to repeat them in either the order presented (forward span) or in reverse order (backwards span). After two consecutive failures of the same length, the test was stopped. Scores were derived as the length of longest correct series for forward and backward recall.

## 3. MSE versus phase-randomised surrogates

To evaluate whether empirical multiscale entropy (MSE) reflected genuine non-linear signal dynamics or linear autocorrelation properties, empirical sample entropy values were compared against phase-randomised surrogate distributions. For each subject and region, 100 surrogate time series were generated by taking the Fourier transform of the empirical signal, preserving the exact amplitude spectrum, and thus the linear autocorrelation, power spectral density, and low-pass filter profile, and replacing the phase angles with random values uniformly distributed on [0,2π]. Sample entropy was calculated across temporal scale factors for both empirical and surrogate time series. A standardised metric was then derived for each scale:

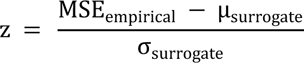

where *μ*_surrogate_ and *σ*_surrogate_ represent the mean and standard deviation of the surrogate distribution, respectively.

Phase randomisation disrupts non-linear phase coupling and temporal asymmetry while preserving linear spectral features, yielding distinct physiological interpretations for dynamic fPET and BOLD fMRI (Fig S1A-D). For fPET, empirical entropy significantly exceeded phase-randomised surrogates at the finest scale (τ = 1, 16 s; z > 0), demonstrating that dynamic glucose utilisation exhibits non-linear ’burstiness’ and high-frequency state transitions that cannot be explained by linear spectral properties alone (Fig. S1G,H). At coarser scales (τ = 2-4), empirical fPET entropy fell below surrogate values (z < 0), consistent with coarse-graining attenuating the non-linear high-frequency structure that distinguishes the empirical signal from its surrogate. Conversely, empirical fMRI entropy fell significantly below surrogate values (z < 0) across all scales (Fig. S1K,L). Rather than indicating a lack of signal complexity, this negative deviation reflects deterministic, non-linearly constrained neurovascular phase coupling [1, 2]. Because phase-randomised surrogates preserve hemodynamic response function (HRF) smoothing, scrambling the underlying phase relationships transforms the smooth, deterministic phase-space trajectory of the BOLD signal into an unconstrained stochastic process (Fig. S1I-L), driving surrogate entropy above empirical levels and confirming that fMRI MSE captures non-linear neurovascular constraints [1].

**Figure S1.**
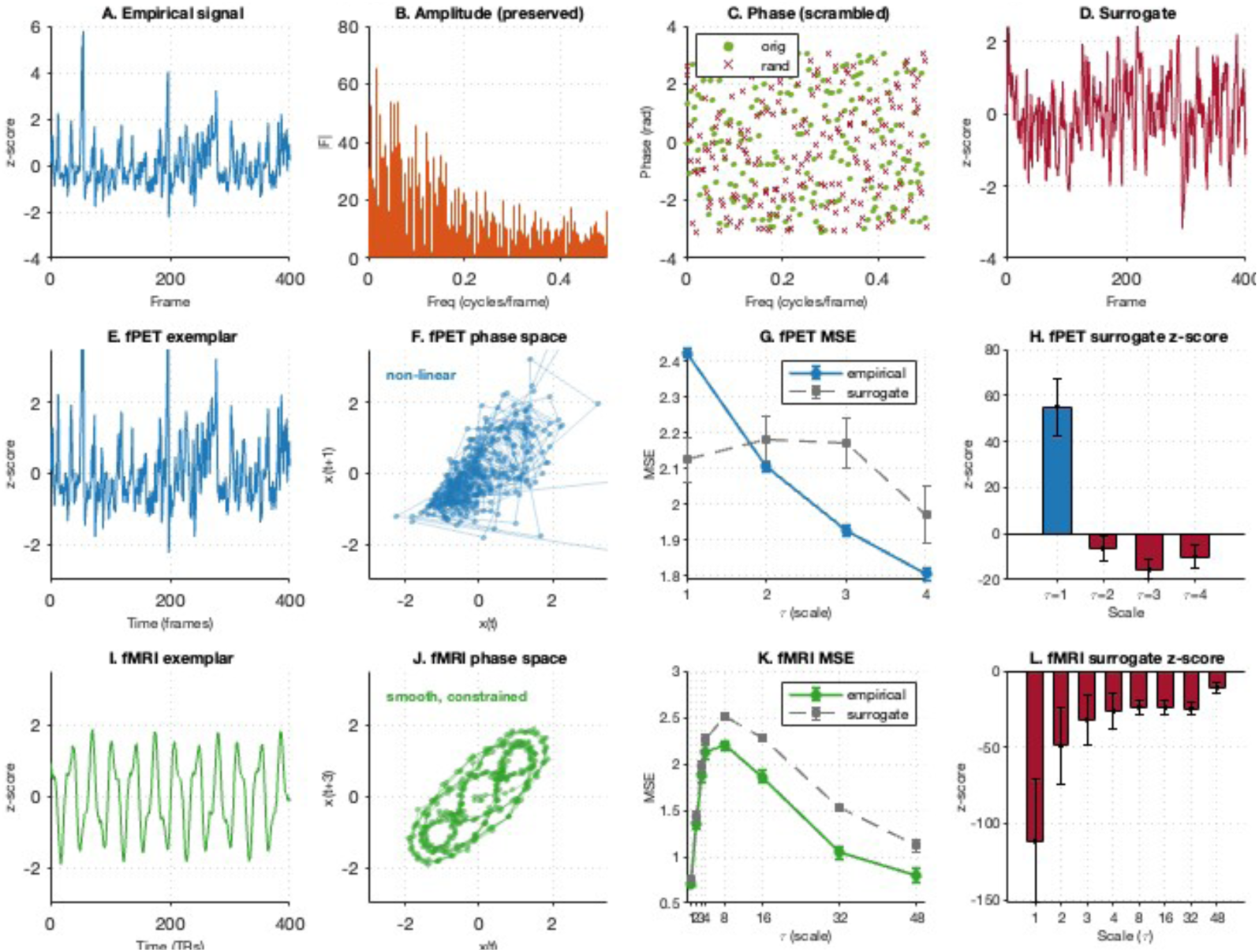
Phase randomisation and scale-dependent multiscale entropy relative to surrogate bounds. (A–D) Schematic of the phase-randomisation procedure. (A) An empirical signal is decomposed by Fourier transform into (B) its amplitude spectrum, which encodes frequency content, and (C) its phase spectrum, which encodes the temporal alignment of frequency components. Phase randomisation preserves the amplitude spectrum but replaces the phase spectrum with random values (open red crosses), yielding (D) a surrogate signal with identical frequency content but no preserved phase structure. (E–H) Dynamic fPET. (E) Illustrative exemplar timeseries with non-linear stochastic dynamics and (F) its two-dimensional phase-space reconstruction (lag τ = 1 frame), shown for dynamical illustration only. (G) Empirical (blue) and phase-randomised surrogate (grey) multiscale entropy (MSE) as a function of scale factor τ (mean ± SD across subjects). (H) Per-scale surrogate z-scores (mean ± SD): empirical fPET MSE exceeds surrogate bounds at τ = 1 (16 s; z > 0, blue) and falls below them at τ = 2–4 (z < 0, red). (I–L) BOLD fMRI. (I) Illustrative exemplar time series with HRF-constrained, smooth dynamics and (J) its phase-space reconstruction (lag τ = 3 TRs), shown for dynamical illustration only. (K) Empirical (green) and surrogate (grey) MSE across τ = 1-48 (mean ± SD across subjects). (L) Per-scale surrogate z-scores: empirical fMRI MSE lies below surrogate bounds at every scale (z < 0 throughout). Panels E, F, I and J are synthetic exemplars generated to illustrate the two dynamical regimes and are not simulations of the empirical data; all MSE and z-score values in G, H, K and L are computed from the empirical datasets. Error bars in G and K are SD across subjects; error bars in H and L are SD of the per-subject z-scores.

## 4. Regional multiscale entropy (MSE)

**Table S2.** Mean and SD regional fPET and fMRI MSE.

|  | fPET MSE |  |  |  |  |  | fMRI MSE |  |  |  |  |  |
|---|---|---|---|---|---|---|---|---|---|---|---|---|
|  | 16s (τ = 1) |  | 32s (τ = 2) |  | 48s (τ = 3) |  | 4s (τ = 4) |  | 8s (τ = 8) |  | 16s (τ = 16) |  |
|  | Mean | SD | Mean | SD | Mean | SD | Mean | SD | Mean | SD | Mean | SD |
| Visual Central: Extra Striate Cortex 1 L | 2.12 | 0.02 | 1.84 | 0.02 | 1.68 | 0.03 | 2.09 | 0.10 | 2.22 | 0.07 | 1.91 | 0.10 |
| Visual Central: Extra Striate Cortex 2 L | 1.90 | 0.02 | 1.65 | 0.02 | 1.51 | 0.02 | 2.04 | 0.12 | 2.18 | 0.10 | 1.92 | 0.13 |
| Visual Central: Striate Cortex 1 L | 1.75 | 0.03 | 1.52 | 0.03 | 1.40 | 0.03 | 2.03 | 0.15 | 2.18 | 0.12 | 1.89 | 0.15 |
| Visual Central: Extra Striate Cortex 3 L | 1.81 | 0.02 | 1.57 | 0.02 | 1.44 | 0.03 | 2.09 | 0.10 | 2.21 | 0.08 | 1.90 | 0.11 |
| Visual Peripheral: Extra Striate Inferior 1 L | 2.16 | 0.01 | 1.87 | 0.02 | 1.71 | 0.03 | 2.13 | 0.09 | 2.19 | 0.08 | 1.86 | 0.11 |
| Visual Peripheral: Striate Cortex Calcarine 1 L | 1.94 | 0.02 | 1.68 | 0.03 | 1.54 | 0.03 | 2.10 | 0.11 | 2.18 | 0.10 | 1.84 | 0.13 |
| Visual Peripheral: Extra Striate CortexSup 1 L | 1.69 | 0.03 | 1.46 | 0.03 | 1.33 | 0.03 | 2.09 | 0.10 | 2.20 | 0.09 | 1.88 | 0.13 |
| Somatomotor A: 1 L | 1.77 | 0.03 | 1.53 | 0.03 | 1.40 | 0.03 | 2.02 | 0.14 | 2.17 | 0.10 | 1.87 | 0.14 |
| Somatomotor A: 2 L | 1.76 | 0.03 | 1.53 | 0.03 | 1.40 | 0.02 | 2.06 | 0.12 | 2.17 | 0.09 | 1.84 | 0.13 |
| Somatomotor B: Auditory 1 L | 1.86 | 0.02 | 1.62 | 0.02 | 1.48 | 0.02 | 2.11 | 0.10 | 2.18 | 0.09 | 1.85 | 0.13 |
| Somatomotor B: S2 1 L | 2.18 | 0.02 | 1.89 | 0.03 | 1.73 | 0.03 | 2.14 | 0.09 | 2.17 | 0.09 | 1.81 | 0.15 |
| Somatomotor B: S2 2 L | 2.12 | 0.02 | 1.84 | 0.03 | 1.68 | 0.03 | 2.11 | 0.08 | 2.20 | 0.08 | 1.85 | 0.13 |
| Somatomotor B: Central 1 L | 1.85 | 0.02 | 1.60 | 0.02 | 1.47 | 0.03 | 2.01 | 0.13 | 2.15 | 0.12 | 1.85 | 0.14 |
| Dorsal Attention A: Temporal Occipital 1 L | 2.13 | 0.01 | 1.84 | 0.02 | 1.69 | 0.02 | 2.11 | 0.08 | 2.21 | 0.06 | 1.90 | 0.09 |
| Dorsal Attention A: Parietal Occipital 1 L | 1.96 | 0.02 | 1.70 | 0.03 | 1.55 | 0.03 | 2.10 | 0.10 | 2.19 | 0.09 | 1.89 | 0.14 |
| Dorsal Attention A: Superior Parietal Lobule 1 L | 1.91 | 0.02 | 1.66 | 0.02 | 1.52 | 0.03 | 2.11 | 0.08 | 2.22 | 0.07 | 1.91 | 0.10 |
| Dorsal Attention B: Post Central 1 L | 1.74 | 0.03 | 1.51 | 0.03 | 1.38 | 0.03 | 2.07 | 0.10 | 2.21 | 0.07 | 1.91 | 0.12 |
| Dorsal Attention B: Post Central 2 L | 1.72 | 0.06 | 1.50 | 0.06 | 1.37 | 0.06 | 2.06 | 0.12 | 2.19 | 0.09 | 1.89 | 0.15 |
| Dorsal Attention B: Post Central 3 L | 1.74 | 0.04 | 1.50 | 0.04 | 1.38 | 0.04 | 2.08 | 0.10 | 2.19 | 0.08 | 1.90 | 0.13 |
| Dorsal Attention B: Frontal Eye Fields 1 L | 1.65 | 0.03 | 1.43 | 0.03 | 1.31 | 0.03 | 2.14 | 0.07 | 2.22 | 0.06 | 1.87 | 0.09 |
| Salience Ventral Attention A: Parietal Operculum 1 L | 1.95 | 0.02 | 1.69 | 0.03 | 1.55 | 0.03 | 2.12 | 0.07 | 2.24 | 0.06 | 1.90 | 0.11 |
| Salience Ventral Attention A: Insula: 1 L | 2.04 | 0.02 | 1.77 | 0.02 | 1.62 | 0.03 | 2.19 | 0.06 | 2.19 | 0.06 | 1.80 | 0.11 |
| Salience Ventral Attention A: Insula: 2 L | 2.27 | 0.02 | 1.96 | 0.03 | 1.80 | 0.03 | 2.18 | 0.06 | 2.21 | 0.07 | 1.82 | 0.11 |
| Salience Ventral Attention A: Parietal Medial 1 L | 1.94 | 0.03 | 1.69 | 0.03 | 1.54 | 0.03 | 2.15 | 0.08 | 2.22 | 0.07 | 1.84 | 0.12 |
| Salience Ventral Attention A: Frontal Medial 1 L | 1.82 | 0.03 | 1.58 | 0.03 | 1.45 | 0.03 | 2.13 | 0.07 | 2.23 | 0.06 | 1.87 | 0.10 |
| Salience Ventral Attention B: Lateral Prefrontal Cortex 1 L | 1.96 | 0.02 | 1.70 | 0.02 | 1.56 | 0.03 | 2.14 | 0.09 | 2.21 | 0.08 | 1.88 | 0.11 |
| Salience Ventral Attention B: Medial Posterior Prefrontal 1 L | 1.84 | 0.03 | 1.59 | 0.03 | 1.45 | 0.03 | 2.18 | 0.07 | 2.21 | 0.06 | 1.82 | 0.10 |
| Limbic A Temporal Pole 1 L | 2.03 | 0.01 | 1.76 | 0.02 | 1.62 | 0.02 | 2.09 | 0.12 | 2.15 | 0.09 | 1.87 | 0.10 |
| Limbic A: Temporal Pole 2 L | 2.09 | 0.01 | 1.82 | 0.02 | 1.66 | 0.02 | 2.12 | 0.11 | 2.14 | 0.09 | 1.83 | 0.10 |
| Limbic B: Orbital Frontal Cortex 1 L | 2.08 | 0.01 | 1.81 | 0.02 | 1.65 | 0.02 | 2.09 | 0.11 | 2.16 | 0.09 | 1.87 | 0.09 |
| Control A: Intraparietal Sulcus 1 L | 1.65 | 0.04 | 1.43 | 0.03 | 1.31 | 0.04 | 2.13 | 0.08 | 2.25 | 0.06 | 1.90 | 0.11 |
| Control A: Lateral Prefrontal Cortex 1 L | 1.77 | 0.02 | 1.54 | 0.03 | 1.41 | 0.03 | 2.14 | 0.08 | 2.22 | 0.06 | 1.88 | 0.10 |
| Control A: Lateral Prefrontal Cortex 2 L | 1.79 | 0.03 | 1.55 | 0.03 | 1.42 | 0.03 | 2.14 | 0.07 | 2.23 | 0.06 | 1.88 | 0.10 |
| Control B: Lateral Prefrontal Cortexv 1 L | 2.01 | 0.02 | 1.74 | 0.02 | 1.60 | 0.02 | 2.06 | 0.13 | 2.15 | 0.12 | 1.87 | 0.13 |
| Control C: Precuneus 1 L | 1.99 | 0.02 | 1.73 | 0.03 | 1.58 | 0.03 | 2.14 | 0.08 | 2.25 | 0.07 | 1.89 | 0.11 |
| Control C: Precuneus 2 L | 1.87 | 0.03 | 1.63 | 0.03 | 1.49 | 0.03 | 2.11 | 0.09 | 2.23 | 0.08 | 1.91 | 0.12 |
| Control C: Cingulate Posterior 1 L | 1.73 | 0.03 | 1.50 | 0.03 | 1.37 | 0.03 | 2.17 | 0.07 | 2.24 | 0.06 | 1.85 | 0.11 |
| Default A: Dorsal Prefrontal Cortex 1 L | 1.91 | 0.03 | 1.66 | 0.03 | 1.52 | 0.03 | 2.15 | 0.07 | 2.24 | 0.06 | 1.90 | 0.10 |
| Default A: Precuneus Posterior Cingulate Cortex1 L | 1.93 | 0.02 | 1.67 | 0.02 | 1.53 | 0.02 | 2.15 | 0.07 | 2.26 | 0.06 | 1.89 | 0.12 |
| Default A: Medial Prefrontal Cortex 1 L | 1.91 | 0.02 | 1.65 | 0.02 | 1.51 | 0.03 | 2.13 | 0.09 | 2.19 | 0.08 | 1.86 | 0.09 |
| Default B: Temp 1 L | 2.06 | 0.01 | 1.78 | 0.02 | 1.63 | 0.02 | 2.12 | 0.09 | 2.20 | 0.07 | 1.90 | 0.10 |
| Default B: Temp 2 L | 2.03 | 0.02 | 1.76 | 0.02 | 1.61 | 0.02 | 2.10 | 0.10 | 2.21 | 0.08 | 1.90 | 0.11 |
| Default B: Inferior Parietal Lobule 1 L | 1.89 | 0.02 | 1.64 | 0.02 | 1.50 | 0.02 | 2.14 | 0.07 | 2.25 | 0.06 | 1.90 | 0.11 |
| Default B: Dorsal Prefrontal Cortex 1 L | 1.98 | 0.02 | 1.72 | 0.02 | 1.57 | 0.02 | 2.13 | 0.08 | 2.22 | 0.07 | 1.90 | 0.10 |
| Default B: Lateral Prefrontal Cortex 1 L | 1.95 | 0.02 | 1.69 | 0.03 | 1.55 | 0.03 | 2.13 | 0.08 | 2.25 | 0.05 | 1.93 | 0.11 |
| Default B: Ventral Prefrontal Cortex 1 L | 2.08 | 0.02 | 1.81 | 0.03 | 1.65 | 0.03 | 2.18 | 0.08 | 2.21 | 0.07 | 1.84 | 0.10 |
| Default B: Ventral Prefrontal Cortex 2 L | 1.99 | 0.02 | 1.73 | 0.02 | 1.58 | 0.02 | 2.14 | 0.08 | 2.22 | 0.06 | 1.89 | 0.09 |
| Default C: RetroSuperior Parietal Lobuleenial 1 L | 2.14 | 0.02 | 1.86 | 0.02 | 1.70 | 0.03 | 2.16 | 0.08 | 2.23 | 0.06 | 1.87 | 0.12 |
| Default C: Parahippocampal Cortex 1 L | 2.22 | 0.02 | 1.93 | 0.02 | 1.76 | 0.03 | 2.20 | 0.06 | 2.20 | 0.06 | 1.82 | 0.08 |
| Temporal Parietal 1 L | 2.15 | 0.02 | 1.87 | 0.02 | 1.71 | 0.03 | 2.11 | 0.11 | 2.20 | 0.09 | 1.89 | 0.12 |
| Visual Central: Extra Striate Cortex 1 R | 2.02 | 0.02 | 1.75 | 0.02 | 1.60 | 0.02 | 2.12 | 0.08 | 2.21 | 0.08 | 1.88 | 0.10 |
| Visual Central: Extra Striate Cortex 2 R | 1.92 | 0.02 | 1.67 | 0.02 | 1.53 | 0.02 | 2.04 | 0.11 | 2.19 | 0.10 | 1.91 | 0.12 |
| Visual Central: Extra Striate Cortex 3 R | 1.88 | 0.02 | 1.63 | 0.02 | 1.49 | 0.03 | 2.08 | 0.08 | 2.21 | 0.07 | 1.92 | 0.12 |
| Visual Peripheral: Striate Cortex Calcarine 1 R | 1.89 | 0.02 | 1.64 | 0.02 | 1.50 | 0.02 | 2.06 | 0.12 | 2.18 | 0.11 | 1.86 | 0.12 |
| Visual Peripheral: Extra Striate Inferior 1 R | 1.94 | 0.02 | 1.68 | 0.02 | 1.53 | 0.02 | 2.12 | 0.09 | 2.17 | 0.09 | 1.82 | 0.12 |
| Visual Peripheral: Extra Striate Superior 1 R | 1.76 | 0.02 | 1.52 | 0.02 | 1.39 | 0.02 | 2.06 | 0.11 | 2.20 | 0.09 | 1.90 | 0.12 |
| Somatomotor A: 1 R | 2.14 | 0.02 | 1.86 | 0.03 | 1.70 | 0.04 | 2.06 | 0.13 | 2.19 | 0.10 | 1.88 | 0.14 |
| Somatomotor A: 2 R | 1.77 | 0.03 | 1.54 | 0.03 | 1.40 | 0.03 | 2.00 | 0.16 | 2.15 | 0.13 | 1.88 | 0.17 |
| Somatomotor A: 3 R | 1.51 | 0.05 | 1.31 | 0.04 | 1.19 | 0.04 | 2.04 | 0.14 | 2.17 | 0.10 | 1.90 | 0.16 |
| Somatomotor A: 4 R | 1.80 | 0.03 | 1.56 | 0.03 | 1.43 | 0.02 | 2.05 | 0.13 | 2.17 | 0.10 | 1.85 | 0.13 |
| Somatomotor B: Auditory 1 R | 1.84 | 0.02 | 1.60 | 0.02 | 1.46 | 0.02 | 2.11 | 0.09 | 2.18 | 0.07 | 1.84 | 0.12 |
| Somatomotor B: S2 1 R | 2.14 | 0.02 | 1.85 | 0.03 | 1.69 | 0.04 | 2.13 | 0.09 | 2.18 | 0.07 | 1.82 | 0.13 |
| Somatomotor B: S2 2 R | 2.01 | 0.04 | 1.75 | 0.04 | 1.59 | 0.04 | 2.07 | 0.11 | 2.19 | 0.07 | 1.89 | 0.14 |
| Somatomotor B: Central 1 R | 1.68 | 0.02 | 1.46 | 0.02 | 1.34 | 0.02 | 1.97 | 0.15 | 2.13 | 0.14 | 1.85 | 0.16 |
| Dorsal Attention A: Temporal Occipital 1 R | 2.02 | 0.01 | 1.75 | 0.02 | 1.60 | 0.02 | 2.08 | 0.09 | 2.19 | 0.08 | 1.91 | 0.11 |
| Dorsal Attention A: Parietal Occipital 1 R | 1.99 | 0.02 | 1.72 | 0.02 | 1.58 | 0.02 | 2.10 | 0.09 | 2.22 | 0.06 | 1.92 | 0.09 |
| Dorsal Attention A: Superior Parietal Lobule 1 R | 1.66 | 0.03 | 1.44 | 0.03 | 1.31 | 0.03 | 2.09 | 0.08 | 2.22 | 0.07 | 1.92 | 0.11 |
| Dorsal Attention B: Post Central 1 R | 1.81 | 0.03 | 1.57 | 0.03 | 1.43 | 0.03 | 2.05 | 0.12 | 2.19 | 0.09 | 1.91 | 0.13 |
| Dorsal Attention B: Post Central 2 R | 1.78 | 0.03 | 1.54 | 0.03 | 1.41 | 0.03 | 2.08 | 0.10 | 2.21 | 0.08 | 1.90 | 0.11 |
| Dorsal Attention B: Frontal Eye Fields 1 R | 1.64 | 0.03 | 1.43 | 0.03 | 1.31 | 0.03 | 2.12 | 0.08 | 2.22 | 0.06 | 1.88 | 0.09 |
| Salience Ventral Attention A: Parietal Operculum 1 R | 2.09 | 0.02 | 1.81 | 0.03 | 1.65 | 0.03 | 2.10 | 0.09 | 2.24 | 0.06 | 1.93 | 0.12 |
| Salience Ventral Attention A: Insula: 1 R | 2.09 | 0.02 | 1.81 | 0.02 | 1.65 | 0.02 | 2.17 | 0.08 | 2.21 | 0.06 | 1.85 | 0.10 |
| Salience Ventral Attention A: Parietal Medial 1 R | 1.93 | 0.02 | 1.68 | 0.03 | 1.53 | 0.03 | 2.15 | 0.08 | 2.22 | 0.07 | 1.85 | 0.10 |
| Salience Ventral Attention A: Frontal Medial 1 R | 1.92 | 0.03 | 1.67 | 0.03 | 1.53 | 0.03 | 2.15 | 0.07 | 2.22 | 0.06 | 1.85 | 0.09 |
| Salience Ventral Attention B: Inferior Parietal Lobule 1 R | 1.75 | 0.03 | 1.52 | 0.03 | 1.39 | 0.03 | 2.12 | 0.08 | 2.26 | 0.06 | 1.94 | 0.11 |
| Salience Ventral Attention B: Lateral Prefrontal Cortex 1 R | 1.85 | 0.02 | 1.61 | 0.02 | 1.48 | 0.02 | 2.12 | 0.10 | 2.21 | 0.09 | 1.90 | 0.11 |
| Salience Ventral Attention B: Medial Posterior Prefrontal 1 R | 1.95 | 0.03 | 1.69 | 0.03 | 1.55 | 0.03 | 2.18 | 0.08 | 2.20 | 0.07 | 1.82 | 0.10 |
| Limbic B: Orbital Frontal Cortex 0 R | 2.11 | 0.01 | 1.84 | 0.02 | 1.68 | 0.02 | 2.08 | 0.12 | 2.14 | 0.09 | 1.87 | 0.11 |
| Limbic A: Temporal Pole 1 R | 2.10 | 0.01 | 1.82 | 0.01 | 1.66 | 0.02 | 2.11 | 0.12 | 2.14 | 0.09 | 1.85 | 0.09 |
| Control A: Intraparietal Sulcus 1 R | 1.49 | 0.05 | 1.30 | 0.04 | 1.19 | 0.05 | 2.11 | 0.09 | 2.23 | 0.06 | 1.92 | 0.11 |
| Control A: Lateral Prefrontal Cortex 1 R | 1.76 | 0.03 | 1.53 | 0.03 | 1.41 | 0.03 | 2.12 | 0.09 | 2.22 | 0.07 | 1.90 | 0.10 |
| Control A: Lateral Prefrontal Cortex 2 R | 1.63 | 0.03 | 1.42 | 0.03 | 1.30 | 0.03 | 2.13 | 0.08 | 2.22 | 0.06 | 1.88 | 0.09 |
| Control B: Temporal 1 R | 2.11 | 0.01 | 1.83 | 0.02 | 1.67 | 0.02 | 2.09 | 0.11 | 2.19 | 0.08 | 1.90 | 0.09 |
| Control B: inferior parietal lobule 1 R | 1.92 | 0.02 | 1.67 | 0.02 | 1.52 | 0.03 | 2.13 | 0.08 | 2.26 | 0.05 | 1.92 | 0.11 |
| Control B: Lateral Prefrontal Cortexd 1 R | 1.96 | 0.02 | 1.70 | 0.03 | 1.56 | 0.03 | 2.13 | 0.08 | 2.25 | 0.06 | 1.91 | 0.10 |
| Control B: Lateral Prefrontal Cortexv 1 R | 1.88 | 0.01 | 1.64 | 0.02 | 1.50 | 0.02 | 2.07 | 0.13 | 2.16 | 0.11 | 1.88 | 0.11 |
| Control C: Cingulate Posterior 1 R | 1.88 | 0.02 | 1.63 | 0.03 | 1.49 | 0.03 | 2.18 | 0.07 | 2.24 | 0.06 | 1.84 | 0.11 |
| Control C: Precuneus 1 R | 2.06 | 0.02 | 1.78 | 0.02 | 1.63 | 0.03 | 2.13 | 0.08 | 2.24 | 0.06 | 1.89 | 0.10 |
| Default A: Inferior Parietal Lobule 1 R | 1.88 | 0.03 | 1.63 | 0.03 | 1.49 | 0.03 | 2.13 | 0.08 | 2.26 | 0.05 | 1.93 | 0.10 |
| Default A: Dorsal Prefrontal Cortex 1 R | 1.86 | 0.03 | 1.62 | 0.03 | 1.48 | 0.03 | 2.15 | 0.08 | 2.23 | 0.07 | 1.88 | 0.10 |
| Default A: Precuneus Posterior Cingulate Cortex 1 R | 1.98 | 0.02 | 1.72 | 0.02 | 1.58 | 0.03 | 2.15 | 0.07 | 2.26 | 0.07 | 1.89 | 0.13 |
| Default A: Medial Prefrontal Cortex 1 R | 1.99 | 0.02 | 1.73 | 0.02 | 1.58 | 0.02 | 2.13 | 0.10 | 2.19 | 0.08 | 1.86 | 0.09 |
| Default B: Dorsal Prefrontal Cortex 1 R | 1.89 | 0.02 | 1.65 | 0.02 | 1.51 | 0.02 | 2.13 | 0.09 | 2.21 | 0.08 | 1.89 | 0.10 |
| Default B: Ventral Prefrontal Cortex 1 R | 1.99 | 0.02 | 1.73 | 0.02 | 1.58 | 0.03 | 2.16 | 0.09 | 2.19 | 0.07 | 1.84 | 0.10 |
| Default B: Ventral Prefrontal Cortex 2 R | 2.08 | 0.02 | 1.81 | 0.02 | 1.66 | 0.03 | 2.14 | 0.08 | 2.22 | 0.07 | 1.88 | 0.10 |
| Default C: RetroSuperior Parietal Lobuleenial 1 R | 2.17 | 0.02 | 1.88 | 0.03 | 1.72 | 0.03 | 2.15 | 0.07 | 2.23 | 0.07 | 1.87 | 0.11 |
| Default C: Parahippocampal Cortex 1 R | 2.25 | 0.01 | 1.94 | 0.02 | 1.77 | 0.02 | 2.16 | 0.07 | 2.19 | 0.06 | 1.85 | 0.09 |
| Temporal Parietal 1 R | 1.98 | 0.02 | 1.72 | 0.02 | 1.57 | 0.02 | 2.11 | 0.11 | 2.17 | 0.09 | 1.87 | 0.12 |
| Temporal Parietal 2 R | 1.92 | 0.02 | 1.67 | 0.02 | 1.53 | 0.02 | 2.09 | 0.11 | 2.20 | 0.09 | 1.90 | 0.11 |
| Temporal Parietal 3 R | 1.77 | 0.02 | 1.53 | 0.03 | 1.40 | 0.03 | 2.10 | 0.10 | 2.22 | 0.08 | 1.90 | 0.13 |

## 5. Multiscale entropy (MSE) and network features

**Table S3.**
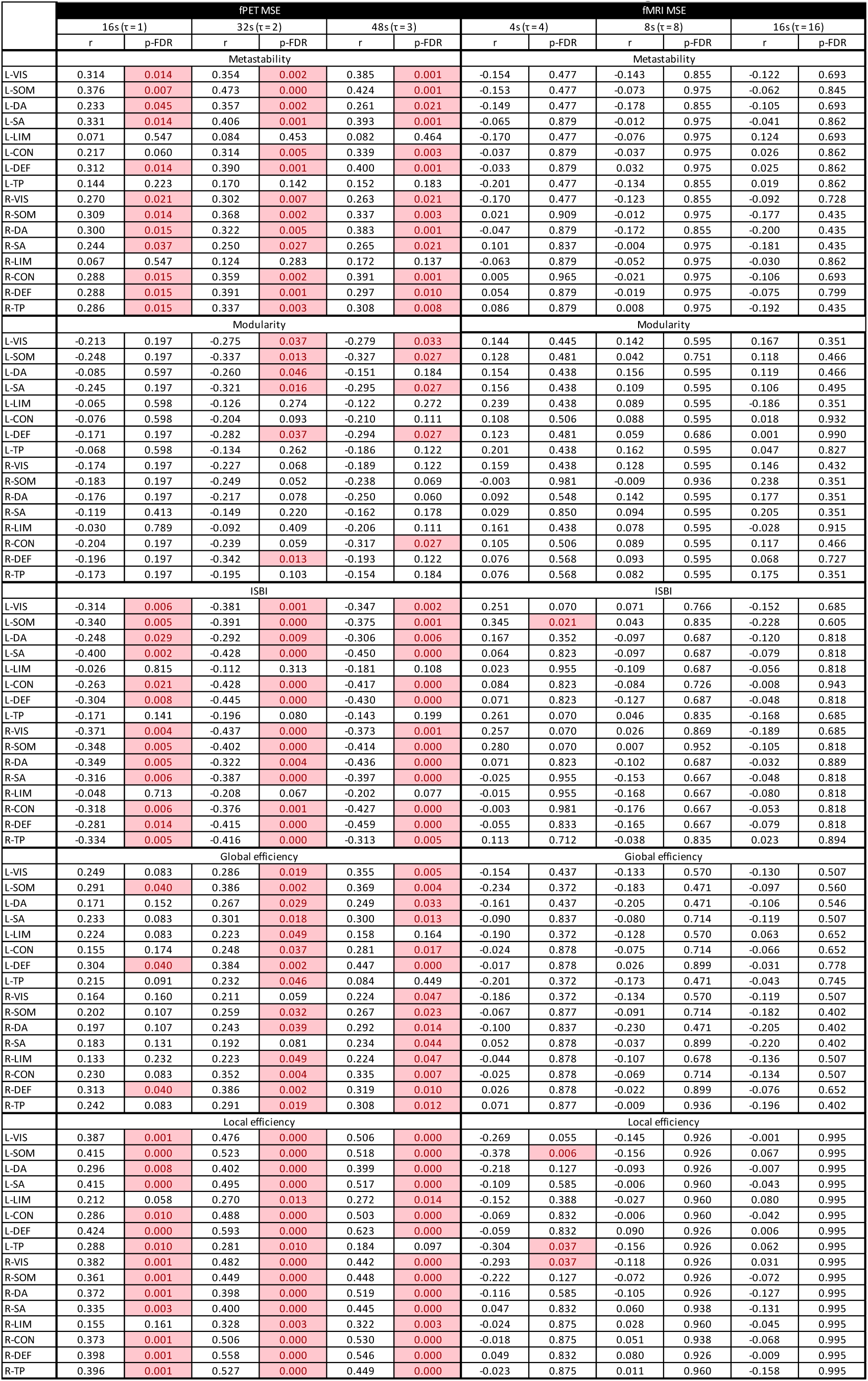
Partial correlation of MSE and network features, controlling head motion.

|  | fPET MSE |  |  |  |  |  | fMRI MSE |  |  |  |  |  |
|---|---|---|---|---|---|---|---|---|---|---|---|---|
|  | 16s (τ = 1) |  | 32s (τ = 2) |  | 48s (τ = 3) |  | 4s (τ = 4) |  | 8s (τ = 8) |  | 16s (τ = 16) |  |
|  | r | p-FDR | r | p-FDR | r | p-FDR | r | p-FDR | r | p-FDR | r | p-FDR |
|  | Metastability |  |  |  |  |  | Metastability |  |  |  |  |  |
| L-VIS | 0.314 | 0.014 | 0.354 | 0.002 | 0.385 | 0.001 | -0.154 | 0.477 | -0.143 | 0.855 | -0.122 | 0.693 |
| L-SOM | 0.376 | 0.007 | 0.473 | 0.000 | 0.424 | 0.001 | -0.153 | 0.477 | -0.073 | 0.975 | -0.062 | 0.845 |
| L-DA | 0.233 | 0.045 | 0.357 | 0.002 | 0.261 | 0.021 | -0.149 | 0.477 | -0.178 | 0.855 | -0.105 | 0.693 |
| L-SA | 0.331 | 0.014 | 0.406 | 0.001 | 0.393 | 0.001 | -0.065 | 0.879 | -0.012 | 0.975 | -0.041 | 0.862 |
| L-LIM | 0.071 | 0.547 | 0.084 | 0.453 | 0.082 | 0.464 | -0.170 | 0.477 | -0.076 | 0.975 | 0.124 | 0.693 |
| L-CON | 0.217 | 0.060 | 0.314 | 0.005 | 0.339 | 0.003 | -0.037 | 0.879 | -0.037 | 0.975 | 0.026 | 0.862 |
| L-DEF | 0.312 | 0.014 | 0.390 | 0.001 | 0.400 | 0.001 | -0.033 | 0.879 | 0.032 | 0.975 | 0.025 | 0.862 |
| L-TP | 0.144 | 0.223 | 0.170 | 0.142 | 0.152 | 0.183 | -0.201 | 0.477 | -0.134 | 0.855 | 0.019 | 0.862 |
| R-VIS | 0.270 | 0.021 | 0.302 | 0.007 | 0.263 | 0.021 | -0.170 | 0.477 | -0.123 | 0.855 | -0.092 | 0.728 |
| R-SOM | 0.309 | 0.014 | 0.368 | 0.002 | 0.337 | 0.003 | 0.021 | 0.909 | -0.012 | 0.975 | -0.177 | 0.435 |
| R-DA | 0.300 | 0.015 | 0.322 | 0.005 | 0.383 | 0.001 | -0.047 | 0.879 | -0.172 | 0.855 | -0.200 | 0.435 |
| R-SA | 0.244 | 0.037 | 0.250 | 0.027 | 0.265 | 0.021 | 0.101 | 0.837 | -0.004 | 0.975 | -0.181 | 0.435 |
| R-LIM | 0.067 | 0.547 | 0.124 | 0.283 | 0.172 | 0.137 | -0.063 | 0.879 | -0.052 | 0.975 | -0.030 | 0.862 |
| R-CON | 0.288 | 0.015 | 0.359 | 0.002 | 0.391 | 0.001 | 0.005 | 0.965 | -0.021 | 0.975 | -0.106 | 0.693 |
| R-DEF | 0.288 | 0.015 | 0.391 | 0.001 | 0.297 | 0.010 | 0.054 | 0.879 | -0.019 | 0.975 | -0.075 | 0.799 |
| R-TP | 0.286 | 0.015 | 0.337 | 0.003 | 0.308 | 0.008 | 0.086 | 0.879 | 0.008 | 0.975 | -0.192 | 0.435 |
|  | Modularity |  |  |  |  |  | Modularity |  |  |  |  |  |
| L-VIS | -0.213 | 0.197 | -0.275 | 0.037 | -0.279 | 0.033 | 0.144 | 0.445 | 0.142 | 0.595 | 0.167 | 0.351 |
| L-SOM | -0.248 | 0.197 | -0.337 | 0.013 | -0.327 | 0.027 | 0.128 | 0.481 | 0.042 | 0.751 | 0.118 | 0.466 |
| L-DA | -0.085 | 0.597 | -0.260 | 0.046 | -0.151 | 0.184 | 0.154 | 0.438 | 0.156 | 0.595 | 0.119 | 0.466 |
| L-SA | -0.245 | 0.197 | -0.321 | 0.016 | -0.295 | 0.027 | 0.156 | 0.438 | 0.109 | 0.595 | 0.106 | 0.495 |
| L-LIM | -0.065 | 0.598 | -0.126 | 0.274 | -0.122 | 0.272 | 0.239 | 0.438 | 0.089 | 0.595 | -0.186 | 0.351 |
| L-CON | -0.076 | 0.598 | -0.204 | 0.093 | -0.210 | 0.111 | 0.108 | 0.506 | 0.088 | 0.595 | 0.018 | 0.932 |
| L-DEF | -0.171 | 0.197 | -0.282 | 0.037 | -0.294 | 0.027 | 0.123 | 0.481 | 0.059 | 0.686 | 0.001 | 0.990 |
| L-TP | -0.068 | 0.598 | -0.134 | 0.262 | -0.186 | 0.122 | 0.201 | 0.438 | 0.162 | 0.595 | 0.047 | 0.827 |
| R-VIS | -0.174 | 0.197 | -0.227 | 0.068 | -0.189 | 0.122 | 0.159 | 0.438 | 0.128 | 0.595 | 0.146 | 0.432 |
| R-SOM | -0.183 | 0.197 | -0.249 | 0.052 | -0.238 | 0.069 | -0.003 | 0.981 | -0.009 | 0.936 | 0.238 | 0.351 |
| R-DA | -0.176 | 0.197 | -0.217 | 0.078 | -0.250 | 0.060 | 0.092 | 0.548 | 0.142 | 0.595 | 0.177 | 0.351 |
| R-SA | -0.119 | 0.413 | -0.149 | 0.220 | -0.162 | 0.178 | 0.029 | 0.850 | 0.094 | 0.595 | 0.205 | 0.351 |
| R-LIM | -0.030 | 0.789 | -0.092 | 0.409 | -0.206 | 0.111 | 0.161 | 0.438 | 0.078 | 0.595 | -0.028 | 0.915 |
| R-CON | -0.204 | 0.197 | -0.239 | 0.059 | -0.317 | 0.027 | 0.105 | 0.506 | 0.089 | 0.595 | 0.117 | 0.466 |
| R-DEF | -0.196 | 0.197 | -0.342 | 0.013 | -0.193 | 0.122 | 0.076 | 0.568 | 0.093 | 0.595 | 0.068 | 0.727 |
| R-TP | -0.173 | 0.197 | -0.195 | 0.103 | -0.154 | 0.184 | 0.076 | 0.568 | 0.082 | 0.595 | 0.175 | 0.351 |
|  | ISBI |  |  |  |  |  | ISBI |  |  |  |  |  |
| L-VIS | -0.314 | 0.006 | -0.381 | 0.001 | -0.347 | 0.002 | 0.251 | 0.070 | 0.071 | 0.766 | -0.152 | 0.685 |
| L-SOM | -0.340 | 0.005 | -0.391 | 0.000 | -0.375 | 0.001 | 0.345 | 0.021 | 0.043 | 0.835 | -0.228 | 0.605 |
| L-DA | -0.248 | 0.029 | -0.292 | 0.009 | -0.306 | 0.006 | 0.167 | 0.352 | -0.097 | 0.687 | -0.120 | 0.818 |
| L-SA | -0.400 | 0.002 | -0.428 | 0.000 | -0.450 | 0.000 | 0.064 | 0.823 | -0.097 | 0.687 | -0.079 | 0.818 |
| L-LIM | -0.026 | 0.815 | -0.112 | 0.313 | -0.181 | 0.108 | 0.023 | 0.955 | -0.109 | 0.687 | -0.056 | 0.818 |
| L-CON | -0.263 | 0.021 | -0.428 | 0.000 | -0.417 | 0.000 | 0.084 | 0.823 | -0.084 | 0.726 | -0.008 | 0.943 |
| L-DEF | -0.304 | 0.008 | -0.445 | 0.000 | -0.430 | 0.000 | 0.071 | 0.823 | -0.127 | 0.687 | -0.048 | 0.818 |
| L-TP | -0.171 | 0.141 | -0.196 | 0.080 | -0.143 | 0.199 | 0.261 | 0.070 | 0.046 | 0.835 | -0.168 | 0.685 |
| R-VIS | -0.371 | 0.004 | -0.437 | 0.000 | -0.373 | 0.001 | 0.257 | 0.070 | 0.026 | 0.869 | -0.189 | 0.685 |
| R-SOM | -0.348 | 0.005 | -0.402 | 0.000 | -0.414 | 0.000 | 0.280 | 0.070 | 0.007 | 0.952 | -0.105 | 0.818 |
| R-DA | -0.349 | 0.005 | -0.322 | 0.004 | -0.436 | 0.000 | 0.071 | 0.823 | -0.102 | 0.687 | -0.032 | 0.889 |
| R-SA | -0.316 | 0.006 | -0.387 | 0.000 | -0.397 | 0.000 | -0.025 | 0.955 | -0.153 | 0.667 | -0.048 | 0.818 |
| R-LIM | -0.048 | 0.713 | -0.208 | 0.067 | -0.202 | 0.077 | -0.015 | 0.955 | -0.168 | 0.667 | -0.080 | 0.818 |
| R-CON | -0.318 | 0.006 | -0.376 | 0.001 | -0.427 | 0.000 | -0.003 | 0.981 | -0.176 | 0.667 | -0.053 | 0.818 |
| R-DEF | -0.281 | 0.014 | -0.415 | 0.000 | -0.459 | 0.000 | -0.055 | 0.833 | -0.165 | 0.667 | -0.079 | 0.818 |
| R-TP | -0.334 | 0.005 | -0.416 | 0.000 | -0.313 | 0.005 | 0.113 | 0.712 | -0.038 | 0.835 | 0.023 | 0.894 |
|  | Global efficiency |  |  |  |  |  | Giobal efficiency |  |  |  |  |  |
| L-VIS | 0.249 | 0.083 | 0.286 | 0.019 | 0.355 | 0.005 | -0.154 | 0.437 | -0.133 | 0.570 | -0.130 | 0.507 |
| L-SOM | 0.291 | 0.040 | 0.386 | 0.002 | 0.369 | 0.004 | -0.234 | 0.372 | -0.183 | 0.471 | -0.097 | 0.560 |
| L-DA | 0.171 | 0.152 | 0.267 | 0.029 | 0.249 | 0.033 | -0.161 | 0.437 | -0.205 | 0.471 | -0.106 | 0.546 |
| L-SA | 0.233 | 0.083 | 0.301 | 0.018 | 0.300 | 0.013 | -0.090 | 0.837 | -0.080 | 0.714 | -0.119 | 0.507 |
| L-LIM | 0.224 | 0.083 | 0.223 | 0.049 | 0.158 | 0.164 | -0.190 | 0.372 | -0.128 | 0.570 | 0.063 | 0.652 |
| L-CON | 0.155 | 0.174 | 0.248 | 0.037 | 0.281 | 0.017 | -0.024 | 0.878 | -0.075 | 0.714 | -0.066 | 0.652 |
| L-DEF | 0.304 | 0.040 | 0.384 | 0.002 | 0.447 | 0.000 | -0.017 | 0.878 | 0.026 | 0.899 | -0.031 | 0.778 |
| L-TP | 0.215 | 0.091 | 0.232 | 0.046 | 0.084 | 0.449 | -0.201 | 0.372 | -0.173 | 0.471 | -0.043 | 0.745 |
| R-VIS | 0.164 | 0.160 | 0.211 | 0.059 | 0.224 | 0.047 | -0.186 | 0.372 | -0.134 | 0.570 | -0.119 | 0.507 |
| R-SOM | 0.202 | 0.107 | 0.259 | 0.032 | 0.267 | 0.023 | -0.067 | 0.877 | -0.091 | 0.714 | -0.182 | 0.402 |
| R-DA | 0.197 | 0.107 | 0.243 | 0.039 | 0.292 | 0.014 | -0.100 | 0.837 | -0.230 | 0.471 | -0.205 | 0.402 |
| R-SA | 0.183 | 0.131 | 0.192 | 0.081 | 0.234 | 0.044 | 0.052 | 0.878 | -0.037 | 0.899 | -0.220 | 0.402 |
| R-LIM | 0.133 | 0.232 | 0.223 | 0.049 | 0.224 | 0.047 | -0.044 | 0.878 | -0.107 | 0.678 | -0.136 | 0.507 |
| R-CON | 0.230 | 0.083 | 0.352 | 0.004 | 0.335 | 0.007 | -0.025 | 0.878 | -0.069 | 0.714 | -0.134 | 0.507 |
| R-DEF | 0.313 | 0.040 | 0.386 | 0.002 | 0.319 | 0.010 | 0.026 | 0.878 | -0.022 | 0.899 | -0.076 | 0.652 |
| R-TP | 0.242 | 0.083 | 0.291 | 0.019 | 0.308 | 0.012 | 0.071 | 0.877 | -0.009 | 0.936 | -0.196 | 0.402 |
|  | Local efficiency |  |  |  |  |  | Local efficiency |  |  |  |  |  |
| L-VIS | 0.387 | 0.001 | 0.476 | 0.000 | 0.506 | 0.000 | -0.269 | 0.055 | -0.145 | 0.926 | -0.001 | 0.995 |
| L-SOM | 0.415 | 0.000 | 0.523 | 0.000 | 0.518 | 0.000 | -0.378 | 0.006 | -0.156 | 0.926 | 0.067 | 0.995 |
| L-DA | 0.296 | 0.008 | 0.402 | 0.000 | 0.399 | 0.000 | -0.218 | 0.127 | -0.093 | 0.926 | -0.007 | 0.995 |
| L-SA | 0.415 | 0.000 | 0.495 | 0.000 | 0.517 | 0.000 | -0.109 | 0.585 | -0.006 | 0.960 | -0.043 | 0.995 |
| L-LIM | 0.212 | 0.058 | 0.270 | 0.013 | 0.272 | 0.014 | -0.152 | 0.388 | -0.027 | 0.960 | 0.080 | 0.995 |
| L-CON | 0.286 | 0.010 | 0.488 | 0.000 | 0.503 | 0.000 | -0.069 | 0.832 | -0.006 | 0.960 | -0.042 | 0.995 |
| L-DEF | 0.424 | 0.000 | 0.593 | 0.000 | 0.623 | 0.000 | -0.059 | 0.832 | 0.090 | 0.926 | 0.006 | 0.995 |
| L-TP | 0.288 | 0.010 | 0.281 | 0.010 | 0.184 | 0.097 | -0.304 | 0.037 | -0.156 | 0.926 | 0.062 | 0.995 |
| R-VIS | 0.382 | 0.001 | 0.482 | 0.000 | 0.442 | 0.000 | -0.293 | 0.037 | -0.118 | 0.926 | 0.031 | 0.995 |
| R-SOM | 0.361 | 0.001 | 0.449 | 0.000 | 0.448 | 0.000 | -0.222 | 0.127 | -0.072 | 0.926 | -0.072 | 0.995 |
| R-DA | 0.372 | 0.001 | 0.398 | 0.000 | 0.519 | 0.000 | -0.116 | 0.585 | -0.105 | 0.926 | -0.127 | 0.995 |
| R-SA | 0.335 | 0.003 | 0.400 | 0.000 | 0.445 | 0.000 | 0.047 | 0.832 | 0.060 | 0.938 | -0.131 | 0.995 |
| R-LIM | 0.155 | 0.161 | 0.328 | 0.003 | 0.322 | 0.003 | -0.024 | 0.875 | 0.028 | 0.960 | -0.045 | 0.995 |
| R-CON | 0.373 | 0.001 | 0.506 | 0.000 | 0.530 | 0.000 | -0.018 | 0.875 | 0.051 | 0.938 | -0.068 | 0.995 |
| R-DEF | 0.398 | 0.001 | 0.558 | 0.000 | 0.546 | 0.000 | 0.049 | 0.832 | 0.080 | 0.926 | -0.009 | 0.995 |
| R-TP | 0.396 | 0.001 | 0.527 | 0.000 | 0.449 | 0.000 | -0.023 | 0.875 | 0.011 | 0.960 | -0.158 | 0.995 |

## 6. Multiscale entropy (MSE) and brain features

**Table S4.**
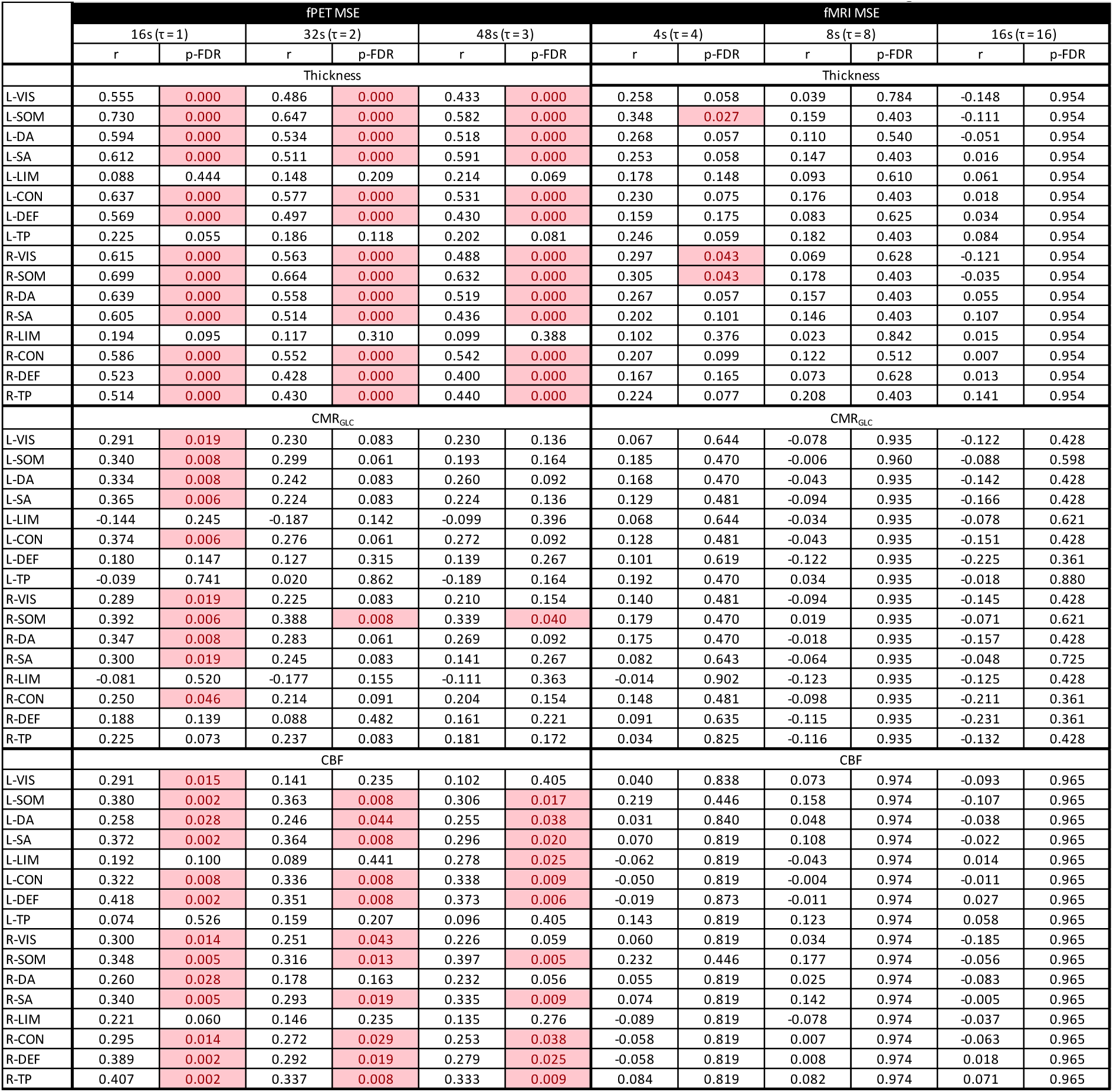
Partial correlation of MSE cortical thickness, CMR_GLC_ and CBF, controlling head motion.

|  | fPET MSE |  |  |  |  |  | fMRI MSE |  |  |  |  |  |
|---|---|---|---|---|---|---|---|---|---|---|---|---|
|  | 16s (τ = 1) |  | 32s (τ = 2) |  | 48s (τ = 3) |  | 4s (τ = 4) |  | 8s (τ = 8) |  | 16s (τ = 16) |  |
|  | r | p-FDR | r | p-FDR | r | p-FDR | r | p-FDR | r | p-FDR | r | p-FDR |
|  | Thickness |  |  |  |  |  | Thickness |  |  |  |  |  |
| L-VIS | 0.555 | 0.000 | 0.486 | 0.000 | 0.433 | 0.000 | 0.258 | 0.058 | 0.039 | 0.784 | -0.148 | 0.954 |
| L-SOM | 0.730 | 0.000 | 0.647 | 0.000 | 0.582 | 0.000 | 0.348 | 0.027 | 0.159 | 0.403 | -0.111 | 0.954 |
| L-DA | 0.594 | 0.000 | 0.534 | 0.000 | 0.518 | 0.000 | 0.268 | 0.057 | 0.110 | 0.540 | -0.051 | 0.954 |
| L-SA | 0.612 | 0.000 | 0.511 | 0.000 | 0.591 | 0.000 | 0.253 | 0.058 | 0.147 | 0.403 | 0.016 | 0.954 |
| L-LIM | 0.088 | 0.444 | 0.148 | 0.209 | 0.214 | 0.069 | 0.178 | 0.148 | 0.093 | 0.610 | 0.061 | 0.954 |
| L-CON | 0.637 | 0.000 | 0.577 | 0.000 | 0.531 | 0.000 | 0.230 | 0.075 | 0.176 | 0.403 | 0.018 | 0.954 |
| L-DEF | 0.569 | 0.000 | 0.497 | 0.000 | 0.430 | 0.000 | 0.159 | 0.175 | 0.083 | 0.625 | 0.034 | 0.954 |
| L-TP | 0.225 | 0.055 | 0.186 | 0.118 | 0.202 | 0.081 | 0.246 | 0.059 | 0.182 | 0.403 | 0.084 | 0.954 |
| R-VIS | 0.615 | 0.000 | 0.563 | 0.000 | 0.488 | 0.000 | 0.297 | 0.043 | 0.069 | 0.628 | -0.121 | 0.954 |
| R-SOM | 0.699 | 0.000 | 0.664 | 0.000 | 0.632 | 0.000 | 0.305 | 0.043 | 0.178 | 0.403 | -0.035 | 0.954 |
| R-DA | 0.639 | 0.000 | 0.558 | 0.000 | 0.519 | 0.000 | 0.267 | 0.057 | 0.157 | 0.403 | 0.055 | 0.954 |
| R-SA | 0.605 | 0.000 | 0.514 | 0.000 | 0.436 | 0.000 | 0.202 | 0.101 | 0.146 | 0.403 | 0.107 | 0.954 |
| R-LIM | 0.194 | 0.095 | 0.117 | 0.310 | 0.099 | 0.388 | 0.102 | 0.376 | 0.023 | 0.842 | 0.015 | 0.954 |
| R-CON | 0.586 | 0.000 | 0.552 | 0.000 | 0.542 | 0.000 | 0.207 | 0.099 | 0.122 | 0.512 | 0.007 | 0.954 |
| R-DEF | 0.523 | 0.000 | 0.428 | 0.000 | 0.400 | 0.000 | 0.167 | 0.165 | 0.073 | 0.628 | 0.013 | 0.954 |
| R-TP | 0.514 | 0.000 | 0.430 | 0.000 | 0.440 | 0.000 | 0.224 | 0.077 | 0.208 | 0.403 | 0.141 | 0.954 |
|  | CMR<sub>GLC</sub> |  |  |  |  |  | CMR<sub>GLC</sub> |  |  |  |  |  |
| L-VIS | 0.291 | 0.019 | 0.230 | 0.083 | 0.230 | 0.136 | 0.067 | 0.644 | -0.078 | 0.935 | -0.122 | 0.428 |
| L-SOM | 0.340 | 0.008 | 0.299 | 0.061 | 0.193 | 0.164 | 0.185 | 0.470 | -0.006 | 0.960 | -0.088 | 0.598 |
| L-DA | 0.334 | 0.008 | 0.242 | 0.083 | 0.260 | 0.092 | 0.168 | 0.470 | -0.043 | 0.935 | -0.142 | 0.428 |
| L-SA | 0.365 | 0.006 | 0.224 | 0.083 | 0.224 | 0.136 | 0.129 | 0.481 | -0.094 | 0.935 | -0.166 | 0.428 |
| L-LIM | -0.144 | 0.245 | -0.187 | 0.142 | -0.099 | 0.396 | 0.068 | 0.644 | -0.034 | 0.935 | -0.078 | 0.621 |
| L-CON | 0.374 | 0.006 | 0.276 | 0.061 | 0.272 | 0.092 | 0.128 | 0.481 | -0.043 | 0.935 | -0.151 | 0.428 |
| L-DEF | 0.180 | 0.147 | 0.127 | 0.315 | 0.139 | 0.267 | 0.101 | 0.619 | -0.122 | 0.935 | -0.225 | 0.361 |
| L-TP | -0.039 | 0.741 | 0.020 | 0.862 | -0.189 | 0.164 | 0.192 | 0.470 | 0.034 | 0.935 | -0.018 | 0.880 |
| R-VIS | 0.289 | 0.019 | 0.225 | 0.083 | 0.210 | 0.154 | 0.140 | 0.481 | -0.094 | 0.935 | -0.145 | 0.428 |
| R-SOM | 0.392 | 0.006 | 0.388 | 0.008 | 0.339 | 0.040 | 0.179 | 0.470 | 0.019 | 0.935 | -0.071 | 0.621 |
| R-DA | 0.347 | 0.008 | 0.283 | 0.061 | 0.269 | 0.092 | 0.175 | 0.470 | -0.018 | 0.935 | -0.157 | 0.428 |
| R-SA | 0.300 | 0.019 | 0.245 | 0.083 | 0.141 | 0.267 | 0.082 | 0.643 | -0.064 | 0.935 | -0.048 | 0.725 |
| R-LIM | -0.081 | 0.520 | -0.177 | 0.155 | -0.111 | 0.363 | -0.014 | 0.902 | -0.123 | 0.935 | -0.125 | 0.428 |
| R-CON | 0.250 | 0.046 | 0.214 | 0.091 | 0.204 | 0.154 | 0.148 | 0.481 | -0.098 | 0.935 | -0.211 | 0.361 |
| R-DEF | 0.188 | 0.139 | 0.088 | 0.482 | 0.161 | 0.221 | 0.091 | 0.635 | -0.115 | 0.935 | -0.231 | 0.361 |
| R-TP | 0.225 | 0.073 | 0.237 | 0.083 | 0.181 | 0.172 | 0.034 | 0.825 | -0.116 | 0.935 | -0.132 | 0.428 |
|  | CBF |  |  |  |  |  | CBF |  |  |  |  |  |
| L-VIS | 0.291 | 0.015 | 0.141 | 0.235 | 0.102 | 0.405 | 0.040 | 0.838 | 0.073 | 0.974 | -0.093 | 0.965 |
| L-SOM | 0.380 | 0.002 | 0.363 | 0.008 | 0.306 | 0.017 | 0.219 | 0.446 | 0.158 | 0.974 | -0.107 | 0.965 |
| L-DA | 0.258 | 0.028 | 0.246 | 0.044 | 0.255 | 0.038 | 0.031 | 0.840 | 0.048 | 0.974 | -0.038 | 0.965 |
| L-SA | 0.372 | 0.002 | 0.364 | 0.008 | 0.296 | 0.020 | 0.070 | 0.819 | 0.108 | 0.974 | -0.022 | 0.965 |
| L-LIM | 0.192 | 0.100 | 0.089 | 0.441 | 0.278 | 0.025 | -0.062 | 0.819 | -0.043 | 0.974 | 0.014 | 0.965 |
| L-CON | 0.322 | 0.008 | 0.336 | 0.008 | 0.338 | 0.009 | -0.050 | 0.819 | -0.004 | 0.974 | -0.011 | 0.965 |
| L-DEF | 0.418 | 0.002 | 0.351 | 0.008 | 0.373 | 0.006 | -0.019 | 0.873 | -0.011 | 0.974 | 0.027 | 0.965 |
| L-TP | 0.074 | 0.526 | 0.159 | 0.207 | 0.096 | 0.405 | 0.143 | 0.819 | 0.123 | 0.974 | 0.058 | 0.965 |
| R-VIS | 0.300 | 0.014 | 0.251 | 0.043 | 0.226 | 0.059 | 0.060 | 0.819 | 0.034 | 0.974 | -0.185 | 0.965 |
| R-SOM | 0.348 | 0.005 | 0.316 | 0.013 | 0.397 | 0.005 | 0.232 | 0.446 | 0.177 | 0.974 | -0.056 | 0.965 |
| R-DA | 0.260 | 0.028 | 0.178 | 0.163 | 0.232 | 0.056 | 0.055 | 0.819 | 0.025 | 0.974 | -0.083 | 0.965 |
| R-SA | 0.340 | 0.005 | 0.293 | 0.019 | 0.335 | 0.009 | 0.074 | 0.819 | 0.142 | 0.974 | -0.005 | 0.965 |
| R-LIM | 0.221 | 0.060 | 0.146 | 0.235 | 0.135 | 0.276 | -0.089 | 0.819 | -0.078 | 0.974 | -0.037 | 0.965 |
| R-CON | 0.295 | 0.014 | 0.272 | 0.029 | 0.253 | 0.038 | -0.058 | 0.819 | 0.007 | 0.974 | -0.063 | 0.965 |
| R-DEF | 0.389 | 0.002 | 0.292 | 0.019 | 0.279 | 0.025 | -0.058 | 0.819 | 0.008 | 0.974 | 0.018 | 0.965 |
| R-TP | 0.407 | 0.002 | 0.337 | 0.008 | 0.333 | 0.009 | 0.084 | 0.819 | 0.082 | 0.974 | 0.071 | 0.965 |

## 7. Age group differences in multiscale entropy (MSE)

**Table S5.**
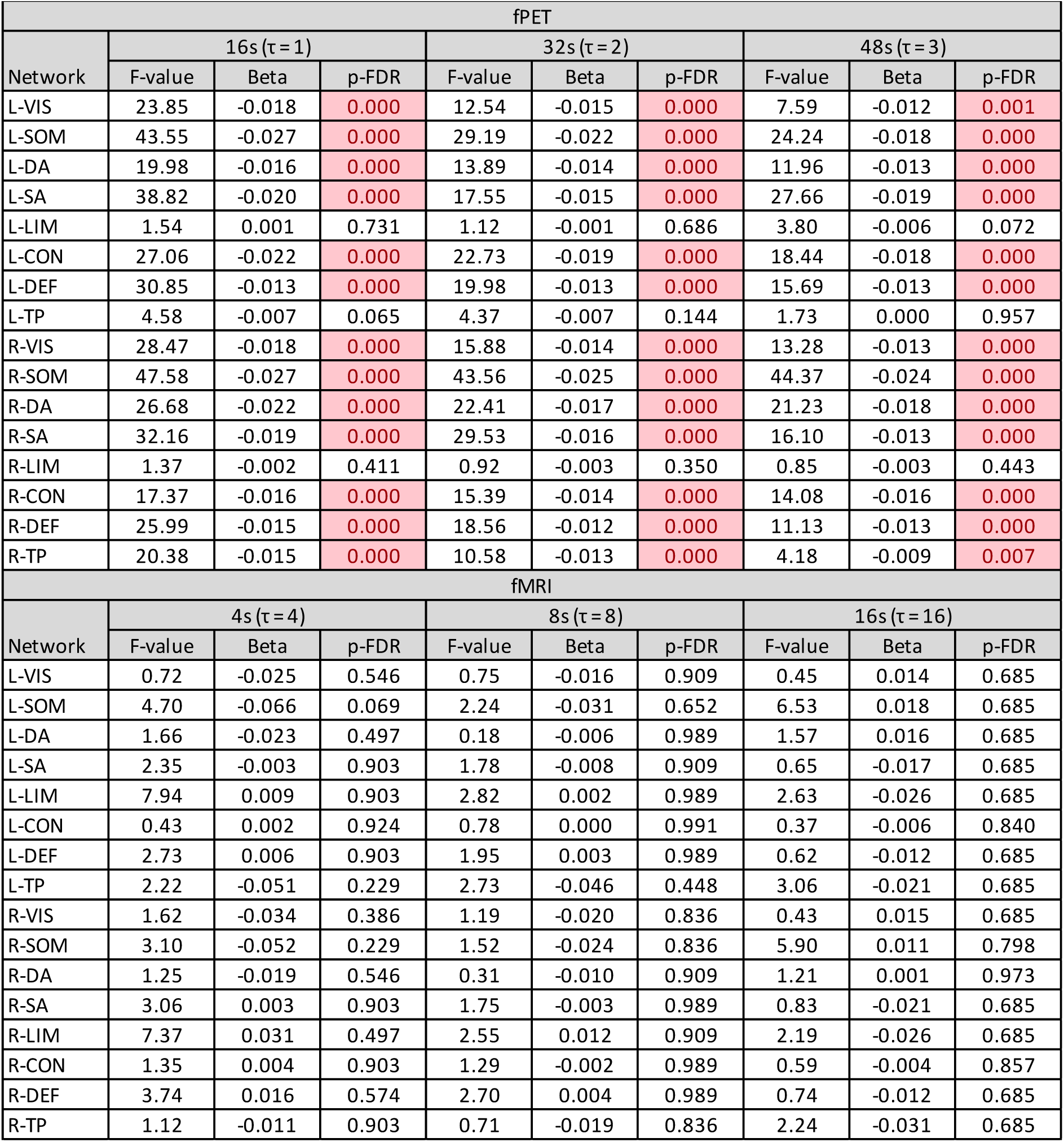
GLMs of age group differences in MSE, controlling head motion. Negative Beta indicates MSE is lower in older vs younger adults.

| fPET |  |  |  |  |  |  |  |  |  |
|---|---|---|---|---|---|---|---|---|---|
| Network | 16s (τ = 1) |  |  | 32s (τ = 2) |  |  | 48s (τ = 3) |  |  |
|  | F-value | Beta | p-FDR | F-value | Beta | p-FDR | F-value | Beta | p-FDR |
| L-VIS | 23.85 | -0.018 | 0.000 | 12.54 | -0.015 | 0.000 | 7.59 | -0.012 | 0.001 |
| L-SOM | 43.55 | -0.027 | 0.000 | 29.19 | -0.022 | 0.000 | 24.24 | -0.018 | 0.000 |
| L-DA | 19.98 | -0.016 | 0.000 | 13.89 | -0.014 | 0.000 | 11.96 | -0.013 | 0.000 |
| L-SA | 38.82 | -0.020 | 0.000 | 17.55 | -0.015 | 0.000 | 27.66 | -0.019 | 0.000 |
| L-LIM | 1.54 | 0.001 | 0.731 | 1.12 | -0.001 | 0.686 | 3.80 | -0.006 | 0.072 |
| L-CON | 27.06 | -0.022 | 0.000 | 22.73 | -0.019 | 0.000 | 18.44 | -0.018 | 0.000 |
| L-DEF | 30.85 | -0.013 | 0.000 | 19.98 | -0.013 | 0.000 | 15.69 | -0.013 | 0.000 |
| L-TP | 4.58 | -0.007 | 0.065 | 4.37 | -0.007 | 0.144 | 1.73 | 0.000 | 0.957 |
| R-VIS | 28.47 | -0.018 | 0.000 | 15.88 | -0.014 | 0.000 | 13.28 | -0.013 | 0.000 |
| R-SOM | 47.58 | -0.027 | 0.000 | 43.56 | -0.025 | 0.000 | 44.37 | -0.024 | 0.000 |
| R-DA | 26.68 | -0.022 | 0.000 | 22.41 | -0.017 | 0.000 | 21.23 | -0.018 | 0.000 |
| R-SA | 32.16 | -0.019 | 0.000 | 29.53 | -0.016 | 0.000 | 16.10 | -0.013 | 0.000 |
| R-LIM | 1.37 | -0.002 | 0.411 | 0.92 | -0.003 | 0.350 | 0.85 | -0.003 | 0.443 |
| R-CON | 17.37 | -0.016 | 0.000 | 15.39 | -0.014 | 0.000 | 14.08 | -0.016 | 0.000 |
| R-DEF | 25.99 | -0.015 | 0.000 | 18.56 | -0.012 | 0.000 | 11.13 | -0.013 | 0.000 |
| R-TP | 20.38 | -0.015 | 0.000 | 10.58 | -0.013 | 0.000 | 4.18 | -0.009 | 0.007 |
| fMRI |  |  |  |  |  |  |  |  |  |
| Network | 4s (τ = 4) |  |  | 8s (τ = 8) |  |  | 16s (τ = 16) |  |  |
|  | F-value | Beta | p-FDR | F-value | Beta | p-FDR | F-value | Beta | p-FDR |
| L-VIS | 0.72 | -0.025 | 0.546 | 0.75 | -0.016 | 0.909 | 0.45 | 0.014 | 0.685 |
| L-SOM | 4.70 | -0.066 | 0.069 | 2.24 | -0.031 | 0.652 | 6.53 | 0.018 | 0.685 |
| L-DA | 1.66 | -0.023 | 0.497 | 0.18 | -0.006 | 0.989 | 1.57 | 0.016 | 0.685 |
| L-SA | 2.35 | -0.003 | 0.903 | 1.78 | -0.008 | 0.909 | 0.65 | -0.017 | 0.685 |
| L-LIM | 7.94 | 0.009 | 0.903 | 2.82 | 0.002 | 0.989 | 2.63 | -0.026 | 0.685 |
| L-CON | 0.43 | 0.002 | 0.924 | 0.78 | 0.000 | 0.991 | 0.37 | -0.006 | 0.840 |
| L-DEF | 2.73 | 0.006 | 0.903 | 1.95 | 0.003 | 0.989 | 0.62 | -0.012 | 0.685 |
| L-TP | 2.22 | -0.051 | 0.229 | 2.73 | -0.046 | 0.448 | 3.06 | -0.021 | 0.685 |
| R-VIS | 1.62 | -0.034 | 0.386 | 1.19 | -0.020 | 0.836 | 0.43 | 0.015 | 0.685 |
| R-SOM | 3.10 | -0.052 | 0.229 | 1.52 | -0.024 | 0.836 | 5.90 | 0.011 | 0.798 |
| R-DA | 1.25 | -0.019 | 0.546 | 0.31 | -0.010 | 0.909 | 1.21 | 0.001 | 0.973 |
| R-SA | 3.06 | 0.003 | 0.903 | 1.75 | -0.003 | 0.989 | 0.83 | -0.021 | 0.685 |
| R-LIM | 7.37 | 0.031 | 0.497 | 2.55 | 0.012 | 0.909 | 2.19 | -0.026 | 0.685 |
| R-CON | 1.35 | 0.004 | 0.903 | 1.29 | -0.002 | 0.989 | 0.59 | -0.004 | 0.857 |
| R-DEF | 3.74 | 0.016 | 0.574 | 2.70 | 0.004 | 0.989 | 0.74 | -0.012 | 0.685 |
| R-TP | 1.12 | -0.011 | 0.903 | 0.71 | -0.019 | 0.836 | 2.24 | -0.031 | 0.685 |

## 8. Significant fPET multiscale entropy (MSE)-cognition clusters

**Table S6.** Significant clusters of MSE and cognition, controlling head motion.

| Analysis | Cognitive domain | MSE (sec) | MSE scale | Size | MNI X | MNI Y | MNI Z | Max T | Mean T | Reg # | Region | % overlap |
|---|---|---|---|---|---|---|---|---|---|---|---|---|
| No Age | Category Switch (RT) | 16 | 1 | 255 | 46 | -13 | 8 | 6.4 | 3.6 | 70 | Central Opercular Cortex Right | 31.8 |
| No Age | Category Switch (RT) | 16 | 1 | 162 | 33 | -21 | -11 | 5.0 | 3.6 | 78 | Hippocampus Right | 40.1 |
| No Age | Category Switch (RT) | 16 | 1 | 146 | 0 | 24 | 36 | 5.4 | 3.5 | 52 | Cingulate Gyrus, anterior division | 32.9 |
| No Age | Category Switch (RT) | 32 | 2 | 212 | 46 | -14 | 9 | 6.7 | 3.6 | 75 | Parietal Operculum Cortex Right | 26.9 |
| No Age | Category Switch (RT) | 32 | 2 | 123 | -43 | -3 | 6 | 4.7 | 3.4 | 14 | Central Opercular Cortex Left | 36.6 |
| No Age | Category Switch (RT) | 32 | 2 | 118 | 33 | -23 | -9 | 5.2 | 3.6 | 78 | Hippocampus Right | 40.7 |
| No Age | Category Switch (RT) | 32 | 2 | 101 | 11 | 12 | 15 | 4.7 | 3.4 | 80 | Caudate Right | 4.0 |
| No Age | Category Switch (RT) | 32 | 2 | 93 | -1 | 29 | 37 | 5.2 | 3.6 | 8 | Paracingulate Gyrus Left | 29.0 |
| No Age | Category Switch (RT) | 32 | 2 | 84 | -9 | 15 | 12 | 4.9 | 3.6 |  | No atlas label | 0.0 |
| No Age | Category Switch (RT) | 48 | 3 | 193 | 46 | -13 | 8 | 5.6 | 3.6 | 70 | Central Opercular Cortex Right | 27.5 |
| No Age | Category Switch (RT) | 48 | 3 | 98 | 33 | -27 | -7 | 5.3 | 3.5 | 78 | Hippocampus Right | 44.9 |
| No Age | Category Switch (RT) | 48 | 3 | 88 | 11 | 9 | 17 | 4.3 | 3.5 | 80 | Caudate Right | 4.5 |
| No Age | Category Switch (RT) | 48 | 3 | 82 | -1 | 28 | 38 | 5.1 | 3.6 | 64 | Paracingulate Gyrus Right | 32.9 |
| No Age | Category Switch (RT) | 48 | 3 | 81 | -9 | 15 | 13 | 4.7 | 3.6 |  | No atlas label | 0.0 |
| No Age | Category Switch (RT) | 48 | 3 | 79 | 1 | 42 | 7 | 5.0 | 3.5 | 52 | Cingulate Gyrus, anterior division | 46.8 |
| No Age | Category Switch (RT) | 48 | 3 | 77 | -4 | -37 | 49 | 5.1 | 3.4 | 54 | Cingulate Gyrus, posterior division | 24.7 |
| No Age | Digit Symbol (Correct) | 16 | 1 | 552 | 0 | -13 | 11 | 6.9 | 3.8 | 23 | Thalamus Left | 18.5 |
| No Age | Digit Symbol (Correct) | 16 | 1 | 219 | 1 | 31 | 29 | 7.1 | 3.7 | 64 | Paracingulate Gyrus Right | 37.9 |
| No Age | Digit Symbol (Correct) | 16 | 1 | 182 | -46 | -12 | 3 | 7.0 | 3.6 | 9 | Insular Cortex Left | 21.4 |
| No Age | Digit Symbol (Correct) | 16 | 1 | 156 | 47 | -11 | 8 | 5.7 | 3.5 | 70 | Central Opercular Cortex Right | 44.2 |
| No Age | Digit Symbol (Correct) | 16 | 1 | 71 | -38 | -52 | 50 | 5.3 | 3.4 | 15 | Superior Parietal Lobule Left | 56.3 |
| No Age | Digit Symbol (Correct) | 16 | 1 | 69 | 19 | -64 | -32 | 6.1 | -3.5 | 128 | Cerebellum 6 Right | 36.2 |
| No Age | Digit Symbol (Correct) | 16 | 1 | 63 | -8 | -63 | -30 | 5.1 | -3.5 | 113 | Cerebellum 8 Left | 47.6 |
| No Age | Digit Symbol (Correct) | 16 | 1 | 60 | -48 | -14 | 47 | 6.3 | 3.2 | 12 | Precentral Gyrus Left | 53.3 |
| No Age | Digit Symbol (Correct) | 16 | 1 | 50 | 47 | -11 | 46 | 5.5 | 3.7 | 68 | Precentral Gyrus Right | 90.0 |
| No Age | Digit Symbol (Correct) | 16 | 1 | 48 | 0 | -82 | -22 | 5.3 | 3.5 | 108 | Cerebellum Crus2 Left | 31.3 |
| No Age | Digit Symbol (Correct) | 16 | 1 | 47 | 46 | 15 | -13 | 5.0 | 3.6 | 84 | Temporal Pole Right | 83.0 |
| No Age | Digit Symbol (Correct) | 16 | 1 | 45 | 28 | 26 | 53 | 4.8 | 3.4 | 61 | Middle Frontal Gyrus Right | 57.8 |
| No Age | Digit Symbol (Correct) | 16 | 1 | 44 | -32 | -37 | -50 | 5.1 | -3.6 | 113 | Cerebellum 8 Left | 40.9 |
| No Age | Digit Symbol (Correct) | 16 | 1 | 44 | 32 | -35 | -4 | 5.9 | 3.9 | 78 | Hippocampus Right | 52.3 |
| No Age | Digit Symbol (Correct) | 32 | 2 | 476 | -1 | -15 | 13 | 6.9 | 3.9 | 23 | Thalamus Left | 22.1 |
| No Age | Digit Symbol (Correct) | 32 | 2 | 211 | 1 | 31 | 30 | 7.4 | 3.7 | 64 | Paracingulate Gyrus Right | 36.5 |
| No Age | Digit Symbol (Correct) | 32 | 2 | 151 | -44 | -8 | 1 | 6.4 | 3.7 | 14 | Central Opercular Cortex Left | 26.5 |
| No Age | Digit Symbol (Correct) | 32 | 2 | 110 | 47 | -13 | 7 | 5.7 | 3.6 | 70 | Central Opercular Cortex Right | 34.5 |
| No Age | Digit Symbol (Correct) | 32 | 2 | 70 | 19 | -65 | -32 | 6.2 | -3.5 | 128 | Cerebellum 6 Right | 37.1 |
| No Age | Digit Symbol (Correct) | 32 | 2 | 65 | -38 | -52 | 50 | 5.8 | 3.5 | 15 | Superior Parietal Lobule Left | 56.9 |
| No Age | Digit Symbol (Correct) | 32 | 2 | 58 | -8 | -63 | -30 | 5.0 | -3.6 | 113 | Cerebellum 8 Left | 48.3 |
| No Age | Digit Symbol (Correct) | 32 | 2 | 58 | -48 | -14 | 46 | 6.2 | 3.3 | 12 | Precentral Gyrus Left | 53.4 |
| No Age | Digit Symbol (Correct) | 32 | 2 | 51 | 47 | -12 | 47 | 5.3 | 3.4 | 68 | Precentral Gyrus Right | 90.2 |
| No Age | Digit Symbol (Correct) | 32 | 2 | 50 | 32 | -35 | -4 | 5.9 | 3.8 | 78 | Hippocampus Right | 56.0 |
| No Age | Digit Symbol (Correct) | 32 | 2 | 47 | 1 | -82 | -22 | 5.5 | 3.5 | 108 | Cerebellum Crus2 Left | 31.9 |
| No Age | Digit Symbol (Correct) | 32 | 2 | 39 | 28 | 28 | 53 | 4.8 | 3.4 | 61 | Middle Frontal Gyrus Right | 46.2 |
| No Age | Digit Symbol (Correct) | 32 | 2 | 38 | -29 | -27 | 63 | 6.1 | 3.8 | 13 | Postcentral Gyrus Left | 55.3 |
| No Age | Digit Symbol (Correct) | 32 | 2 | 37 | 46 | 13 | -13 | 4.9 | 3.5 | 84 | Temporal Pole Right | 91.9 |
| No Age | Digit Symbol (Correct) | 32 | 2 | 36 | -26 | 16 | 57 | 4.6 | 3.5 | 4 | Superior Frontal Gyrus Left | 63.9 |
| No Age | Digit Symbol (Correct) | 32 | 2 | 29 | 32 | -80 | 35 | 4.9 | 3.6 | 104 | Lateral Occipital Cortex, superior division Right | 100.0 |
| No Age | Digit Symbol (Correct) | 32 | 2 | 28 | -49 | -30 | 14 | 4.6 | 3.5 | 19 | Parietal Operculum Cortex Left | 64.3 |
| No Age | Digit Symbol (Correct) | 32 | 2 | 27 | 24 | -90 | -32 | 4.4 | 3.4 | 125 | Cerebellum Crus2 Right | 18.5 |
| No Age | Digit Symbol (Correct) | 32 | 2 | 27 | 8 | -38 | 54 | 4.6 | 3.3 | 69 | Postcentral Gyrus Right | 48.1 |
| No Age | Digit Symbol (Correct) | 32 | 2 | 26 | -46 | -31 | 46 | 4.1 | 3.4 | 13 | Postcentral Gyrus Left | 80.8 |
| No Age | Digit Symbol (Correct) | 48 | 3 | 522 | 0 | -15 | 11 | 6.5 | 3.8 | 23 | Thalamus Left | 19.3 |
| No Age | Digit Symbol (Correct) | 48 | 3 | 195 | 1 | 29 | 30 | 7.0 | 3.7 | 52 | Cingulate Gyrus, anterior division | 37.9 |
| No Age | Digit Symbol (Correct) | 48 | 3 | 152 | -44 | -7 | 0 | 6.8 | 3.6 | 9 | Insular Cortex Left | 28.9 |
| No Age | Digit Symbol (Correct) | 48 | 3 | 108 | 45 | -13 | 7 | 6.0 | 3.5 | 70 | Central Opercular Cortex Right | 30.6 |
| No Age | Digit Symbol (Correct) | 48 | 3 | 56 | -37 | -51 | 50 | 5.6 | 3.5 | 15 | Superior Parietal Lobule Left | 69.6 |
| No Age | Digit Symbol (Correct) | 48 | 3 | 51 | 19 | -65 | -33 | 5.6 | -3.6 | 128 | Cerebellum 6 Right | 39.2 |
| No Age | Digit Symbol (Correct) | 48 | 3 | 51 | 1 | -82 | -22 | 5.8 | 3.6 | 108 | Cerebellum Crus2 Left | 27.5 |
| No Age | Digit Symbol (Correct) | 48 | 3 | 51 | -8 | -62 | -30 | 5.1 | -3.5 | 113 | Cerebellum 8 Left | 43.1 |
| No Age | Digit Symbol (Correct) | 48 | 3 | 50 | -48 | -14 | 47 | 6.1 | 3.6 | 12 | Precentral Gyrus Left | 54.0 |
| No Age | Digit Symbol (Correct) | 48 | 3 | 43 | 32 | -35 | -4 | 6.0 | 3.8 | 78 | Hippocampus Right | 46.5 |
| No Age | Digit Symbol (Correct) | 48 | 3 | 39 | 28 | 26 | 54 | 4.5 | 3.4 | 61 | Middle Frontal Gyrus Right | 48.7 |
| No Age | Digit Symbol (Correct) | 48 | 3 | 36 | 49 | 12 | -10 | 5.2 | 3.6 | 84 | Temporal Pole Right | 91.7 |
| No Age | Digit Symbol (Correct) | 48 | 3 | 36 | -29 | -27 | 63 | 6.1 | 3.8 | 13 | Postcentral Gyrus Left | 58.3 |
| No Age | Digit Symbol (Correct) | 48 | 3 | 30 | 46 | -12 | 43 | 5.0 | 3.4 | 68 | Precentral Gyrus Right | 83.3 |
| No Age | Digit Symbol (Correct) | 48 | 3 | 30 | 29 | -1 | 62 | 4.6 | 3.3 | 60 | Superior Frontal Gyrus Right | 66.7 |
| No Age | Digit Symbol (Correct) | 48 | 3 | 30 | 37 | -25 | 62 | 4.3 | 2.0 | 69 | Postcentral Gyrus Right | 76.7 |
| No Age | Digit Symbol (Correct) | 48 | 3 | 28 | -37 | -39 | -52 | 5.7 | -3.7 | 113 | Cerebellum 8 Left | 42.9 |
| No Age | Digit Symbol (Correct) | 48 | 3 | 28 | 32 | -80 | 36 | 4.9 | 3.5 | 104 | Lateral Occipital Cortex, superior division Right | 100.0 |
| No Age | Digit Symbol (Correct) | 48 | 3 | 27 | -26 | 17 | 55 | 4.5 | 3.5 | 4 | Superior Frontal Gyrus Left | 59.3 |
| No Age | Digit Symbol (Correct) | 48 | 3 | 26 | -38 | 21 | -10 | 4.8 | 3.5 | 1 | Frontal Orbital Cortex Left | 92.3 |
| No Age | Digit Symbol (Correct) | 48 | 3 | 26 | -38 | 21 | -10 | 4.8 | 3.5 | 1 | Frontal Pole | 92.3 |
| No Age | HVLT (Delayed Recall) | 48 | 3 | 32 | -10 | 17 | 13 | 3.9 | 3.2 |  | No atlas label | 0.0 |
| No Age | Stop Signal (RT) | 32 | 2 | 36 | -14 | -81 | -49 | 4.4 | 3.3 | 108 | Region 108 | 2.8 |

Note: Regions are from the Harvard Oxford atlas.

## 9. Significant fMRI multiscale entropy (MSE)-cognition clusters

**Table S7.** Significant clusters of MSE and cognition, controlling head motion and age.

| Analysis | Cognitive_Variable | MSE (Sec) | MSE Scale | Size | MNI X | MNI Y | MNI Z | Max T | Mean T | Reg # | Region | % overlap |
|---|---|---|---|---|---|---|---|---|---|---|---|---|
| No Age | Category Switch (RT) | 4 | 4 | 627 | 42 | -21 | 57 | 5.2 | 3.4 | 69 | Postcentral Gyrus Right | 53.6 |
| No Age | Category Switch (RT) | 8 | 8 | 75 | -41 | -20 | 62 | 3.9 | 3.2 | 13 | Postcentral Gyrus Left | 56.0 |
| No Age | Category Switch (RT) | 8 | 8 | 72 | 34 | -29 | 59 | 5.4 | 3.6 | 69 | Postcentral Gyrus Right | 69.4 |
| No Age | Category Switch (RT) | 8 | 8 | 68 | 43 | -17 | 58 | 4.9 | 3.5 | 68 | Precentral Gyrus Right | 69.1 |
| No Age | Category Switch (RT) | 8 | 8 | 64 | -3 | -23 | 59 | 4.7 | 3.3 | 12 | Precentral Gyrus Left | 59.4 |
| No Age | Category Switch (RT) | 8 | 8 | 59 | -58 | -8 | 36 | 4.0 | 3.3 | 13 | Postcentral Gyrus Left | 52.5 |
| No Age | Digit Symbol (Correct) | 4 | 4 | 997 | -44 | -21 | 52 | 5.5 | 3.4 | 13 | Postcentral Gyrus Left | 64.4 |
| No Age | Digit Symbol (Correct) | 4 | 4 | 372 | 0 | -22 | 63 | 4.5 | 3.3 | 12 | Precentral Gyrus Left | 30.1 |
| No Age | HVLT (Delayed Recall) | 4 | 4 | 4991 | -2 | -23 | 56 | 6.6 | 3.5 | 68 | Precentral Gyrus Right | 23.3 |
| No Age | HVLT (Delayed Recall) | 8 | 8 | 113 | 37 | -30 | 54 | 5.0 | 3.4 | 69 | Postcentral Gyrus Right | 86.7 |
| No Age | HVLT (Delayed Recall) | 8 | 8 | 51 | 37 | -18 | 61 | 4.5 | 3.2 | 68 | Precentral Gyrus Right | 88.2 |
| No Age | Stop Signal (RT) | 4 | 4 | 641 | 3 | -33 | 64 | 5.1 | 3.3 | 68 | Precentral Gyrus Right | 24.2 |
| No Age | Stop Signal (RT) | 4 | 4 | 309 | 33 | -26 | 57 | 4.6 | 3.3 | 68 | Precentral Gyrus Right | 42.4 |
| With Age | Category Switch (RT) | 8 | 8 | 35 | 40 | -17 | 58 | 4.8 | 3.5 | 68 | Precentral Gyrus Right | 100.0 |
| With Age | HVLT (Delayed Recall) | 4 | 4 | 1039 | 22 | -23 | 60 | 5.4 | 3.3 | 68 | Precentral Gyrus Right | 43.6 |
| With Age | HVLT (Delayed Recall) | 4 | 4 | 454 | -29 | -25 | 57 | 5.8 | 3.4 | 13 | Postcentral Gyrus Left | 47.8 |
| With Age | Stop Signal (RT) | 4 | 4 | 229 | 5 | -31 | 64 | 4.5 | 3.3 | 68 | Precentral Gyrus Right | 32.8 |

Note: Regions are from the Harvard Oxford atlas.

## 10. Sensitivity analyses

To validate the robustness of our multiscale entropy (MSE) findings and explore alternative measures of signal complexity, we conducted a comprehensive sensitivity analysis using four distinct entropy measures computed at the native temporal resolution for both fPET (16s) and fMRI (1s). These measures were selected to capture complementary aspects of signal irregularity and complexity, each providing a unique window into the temporal structure of metabolic and hemodynamic signals.

**Sample Entropy** measures the conditional probability that two sequences of length m that are similar will remain similar when extended by one sample. It quantifies the regularity and predictability of time series, with higher values indicating greater irregularity. We computed Sample Entropy with the standard parameters (m = 2, r = 0.15; main manuscript) and also performed a sensitivity analysis with a more liberal tolerance threshold (r = 0.30) to assess the stability of our findings across parameter choices.

**Shannon Entropy** is a measure of information content or uncertainty based on the probability distribution of signal values. Unlike Sample Entropy, Shannon Entropy is insensitive to the temporal ordering of data points, capturing instead the spread and diversity of signal amplitudes. This measure provides a complementary perspective on signal complexity by focusing on the distributional properties rather than the sequential structure of the time series. For fMRI, where the hemodynamic response introduces temporal smoothing, Shannon Entropy may capture different aspects of signal variability compared to Sample Entropy.

**Fuzzy Entropy** extends Sample Entropy by employing a fuzzy membership function to determine pattern similarity rather than a hard threshold. This approach makes the measure more robust to noise and small fluctuations in the signal, as it assigns continuous similarity values rather than binary matches. The exponential similarity function used in Fuzzy Entropy reduces sensitivity to parameter choices and provides more stable estimates for short time series.

**Brain Entropy (BEN)** is defined as Sample Entropy computed on z-scored time series. By standardising each voxel’s time series to zero mean and unit variance, BEN isolates the temporal irregularity of the signal’s fluctuation patterns independent of amplitude differences. This measure is particularly useful for comparing complexity across regions with different baseline signal amplitudes, as it focuses on the relative pattern of fluctuations rather than absolute signal variance.

These four entropy measures were chosen to complement our primary MSE analysis (which focuses on Sample Entropy at multiple scales) by providing additional insights into the complexity of metabolic and hemodynamic signals. While MSE captures how signal irregularity changes across temporal scales, Shannon, Fuzzy, and Brain Entropy each provide unique perspectives on signal complexity that may be differentially sensitive to the properties of fPET and fMRI signals. This multi-measure approach allowed us to assess whether the associations observed with MSE are specific to Sample Entropy or generalise across different entropy formulations.

## Results of Sensitivity Analyses

### Surrogate testing

Surrogate testing revealed that all entropy measures showed highly significant deviations from phase-randomized surrogates (all p < 0.001), confirming that the empirical maps contain systematic information beyond random noise. The negative Cohen’s d values for fPET (ranging from -6,4 to -6.7) and fMRI (−1.33 to -3.3) indicate that the empirical metabolic signal is more regular than phase-randomised surrogates. The fact that 100% of participants showed significant deviations (p < 0.001) across all four entropy measures provides strong validation that these metrics capture genuine signal properties rather than statistical artifacts.

### Regional descriptive analysis

fPET (16 s TR). Regional entropy values were tightly distributed across the 100 parcels for all four measures. Shannon entropy ranged approximately 3.83–4.60 bits (mean ≈ 4.21), Fuzzy entropy ≈ 2.13– 2.55 (mean ≈ 2.32), BEN ≈ 1.88–2.26 (mean ≈ 2.07), and SampEn_r0.3 ≈ 1.34–1.60 (mean ≈ 1.47). The coefficient of variation across regions was small (≈ 3–5%), indicating that at 16 s temporal resolution the entropy of the FDG signal is relatively homogeneous across cortex.

fMRI (1 s TR). Regional entropy values were more variable and, for the tolerance-based measures, substantially lower. Shannon entropy ranged ≈ 4.66–5.02 bits (mean ≈ 4.94), which is higher than fPET. In contrast, Fuzzy entropy ≈ 0.84–1.00 (mean ≈ 0.91), BEN ≈ 0.64–0.73 (mean ≈ 0.70), and Sample Entropy 0.3 ≈ 0.48–0.56 (mean ≈ 0.53) were all markedly lower than their fPET counterparts, roughly 40-70% of the fPET values depending on measure.

The cross-modality offset is a predictable consequence of three factors, none of which reflect a biological difference in “complexity” per se: Temporal sampling rate (TR). fPET frames integrate metabolic activity over 16 s, imposing a strong low-pass filter on the signal. fMRI at 1 s TR retains high-frequency content, including cardiac and respiratory fluctuations and their aliases. This 16-fold difference in temporal resolution produces different autocorrelation structures and amplitude distributions, which directly alter entropy estimates.

Tolerance parameter scaling. For Fuzzy entropy, BEN, and Sample Entropy, the tolerance is set relative to each timeseries’ own standard deviation (r = r-ratio × SD). fMRI timeseries have substantially larger sample-to-sample variance than smoothed fPET timeseries; consequently the absolute tolerance r is larger for fMRI. Because Sample and Fuzzy entropy decrease monotonically as r increases above the noise floor, the fMRI values are systematically lower. This is a well-established property of these estimators and is independent of the underlying physiology.

Shannon entropy behaves in the opposite direction. Shannon entropy depends on the breadth of the amplitude histogram rather than on template-match counts. fMRI’s broader amplitude distribution (more high-frequency content, more physiological noise) populates more histogram bins, yielding higher Shannon entropy than fPET. This divergence between Shannon and the tolerance-based measures is expected and has been reported previously when comparing signals of different spectral content.

In short, the direction of the fPET–fMRI difference is measure-specific: Shannon entropy is higher in fMRI, whereas Fuzzy, BEN, and Sample Entropy are higher in fPET. This pattern is consistent with the estimators’ known sensitivity to sampling rate and tolerance scaling rather than with a genuine difference in neural complexity.

### Entropy Associations with Network Measures, Neurobiology and Cognition

Despite the cross-modality offset in absolute values, the sensitivity analyses revealed highly consistent patterns across all four entropy measures, strongly supporting our main findings from the MSE analysis (Table S7). The relationship between entropy and network dynamics was highly consistent across fPET entropy measures. All four fPET measures showed significant positive correlations with metastability (r = 0.29-0.32) and local efficiency (r = 0.32-0.38), and negative correlations with the integration-segregation balance index (ISBI, r = -0.36 to -0.40). In contrast, fMRI entropy showed no consistent network associations, with the exception of fMRI Shannon Entropy showing some significant correlations (metastability: r = 0.37, p = 0.003; local efficiency: r = 0.47, p < 0.001). This suggests that Shannon Entropy may capture network-relevant information in fMRI that is not captured by Sample Entropy, possibly due to its sensitivity to the broader distributional properties of the BOLD signal.

The neurobiological associations were highly consistent for fPET entropy measures. All four fPET measures showed robust positive correlations with cortical thickness (r = 0.78-0.79, all p < 0.001), CMR_GLC_ (r = 0.50-0.54, all p < 0.001), and CBF (r = 0.40, all p < 0.001). In contrast, no significant neurobiological associations were observed for any fMRI entropy measure (all p > 0.50). This pattern highlights the unique sensitivity of fPET entropy to structural and metabolic brain properties, which is absent in the hemodynamic domain.

All fPET entropy measures also showed age group differences, with older adults exhibiting significantly lower entropy values compared to younger adults (t-values ranging from -9.36 to -9.53, all p < 0.001). For fMRI, only Shannon Entropy showed a significant age group difference, and in the opposite direction (t = 2.73, p = 0.012), with older adults showing higher entropy. This suggests that while hemodynamic signal irregularity (as captured by Shannon Entropy) may increase with age, the metabolic signal complexity (captured by all measures) consistently declines.

For fPET, all measures showed robust positive associations with cognitive performance, particularly with processing speed (Digit Symbol Substitution: r = 0.61-0.62, all p < 0.001), memory (HVLT: r = 0.36-0.38, all p < 0.001), and task switching latency (Category Switch: r =-0.42 to 0.43, all p < 0.001). These associations remained significant after controlling for head motion but were largely attenuated after additionally controlling for age, consistent with the age-related nature of fPET entropy changes. In contrast, none of the fMRI entropy measures showed significant associations with any cognitive measure across any of the four entropy formulations, replicating the null findings from our primary MSE analysis. This dissociation between fPET and fMRI was observed across all entropy measures and cognitive domains, underscoring the fundamental differences in the information captured by metabolic versus hemodynamic signals.

## Consistency with Main Findings

Collectively, these sensitivity analyses provide convergent evidence that our main findings are robust to the choice of entropy measure and parameter settings. The consistent pattern of results across Sample Entropy (r = 0.15 and r = 0.30), Shannon Entropy, Fuzzy Entropy, and BEN demonstrates that:

1. fPET entropy is a robust biomarker of brain ageing, cognition, and neurobiology, with effects consistent across multiple entropy formulations
2. fMRI entropy shows minimal sensitivity to these same individual differences, with only Shannon Entropy showing some network associations and a reversed age effect
3. The dissociation between fPET and fMRI is not specific to Sample Entropy but generalises across complementary measures of signal complexity

The robustness of these findings across four distinct entropy measures, each capturing different aspects of signal irregularity, supports our interpretation that metabolic signal complexity provides unique information about brain function that is not captured by hemodynamic measures. Metabolic signal complexity is tightly linked to the physiology of the brain, including the brain structure, metabolic budget and blood flow; whereas fMRI signal entropy is not. This suggests that complexity in fMRI is not particularly relevant to understanding brain function. In contrast, these sensitivity analyses further validate the biological meaningfulness of fPET entropy as a marker of brain health and cognitive function in ageing.

**Figure S2.**
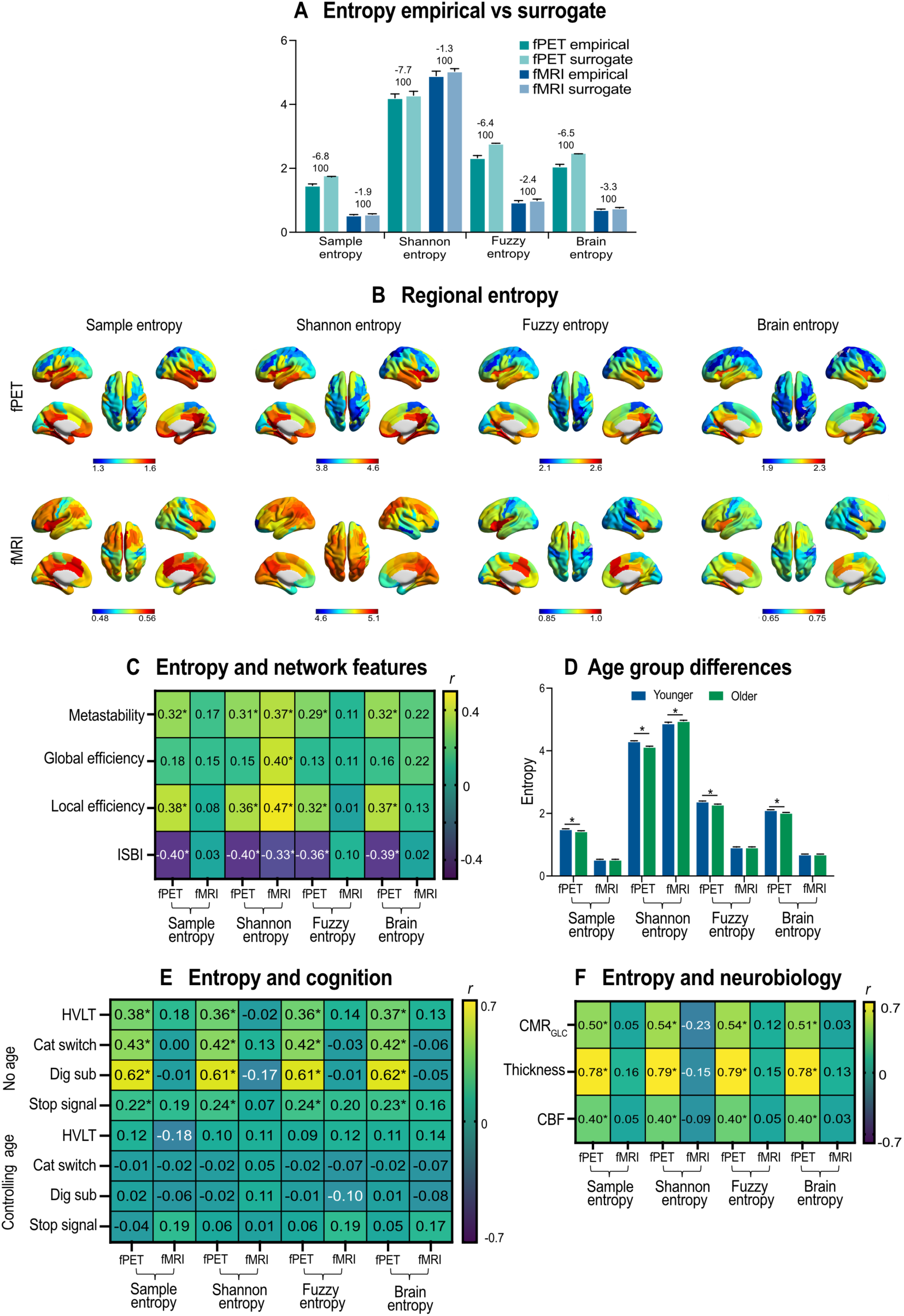
Sensitivity analyses repeating analyses from main manuscript at native fPET frame (16s) and fMRI volume (1s) rates, using different imbedding function (r = 0.30) and entropy measures. (A) empirical versus surrogate null. Values above each pair indicate Cohen’s d and below the percentage of subjects with |z| > 1.96 (i.e., empirical MSE falling outside the surrogate distribution) (B) regional entropy projected to brain surface; (c) entropy and network feature associations; (D) age group differences in entropy; (E) entropy and cognition association; and (F) entropy and neurobiology feature associations. All values are partial correlation (r) controlling for motion, except test of age group differences, which is t-value from GLM of younger vs older age groups. Asterisk (*) indicates significant at p-FDR < 0.05.

